# Preclinical Development and Safety of Laterally Spreading Dual AAV.SPR-*MYO7A* Vectors for the Treatment of Usher Syndrome Type 1B

**DOI:** 10.64898/2026.10.24.760972

**Authors:** Kaitlyn R. Calabro, Eva Andres-Mateos, Christopher Kozlowski, Paige S. Nemec, Emily M. Xu, Sean M. Crosson, Diego Fajardo, Hannah L. Friend, James J. Peterson, Wei Li, Hangning Zhang, H. Lee Sweeney, Victoria Stevenson, Howard Rind, Matthew Leahy, Ryan F. Boyd, Sanford L. Boye, Shannon E. Boye

## Abstract

Usher syndrome type 1B (USH1B), caused by mutations in *MYO7A*, is the most severe form of Usher syndrome and accounts for more than 50% of USH1 cases. Affected patients are born profoundly deaf with vestibular dysfunction and develop progressive retinitis pigmentosa, for which no approved treatment exists. The *MYO7A* coding sequence exceeds the packaging capacity of adeno-associated virus (AAV), the gold standard vector for retinal gene therapy. To overcome this, we developed dual AAV hybrid and overlap platforms to reconstitute full-length MYO7A expression in photoreceptors and retinal pigment epithelium. Modification of cDNA split points, and reduction of non-coding sequence yielded optimized vector pairs that produced higher full-length MYO7A levels with substantially reduced truncated protein. Packaging in the laterally spreading AAV.SPR capsid enabled supraphysiological MYO7A expression in cynomolgus macaques at macular and distal retinal sites without requiring foveal detachment, an important safety consideration for USH1B patients with remaining central vision. A six-month GLP toxicology study confirmed that ATSN-301, an AAV.SPR-based dual vector reconstituting *MYO7A* driven by the smCBA promoter, was well tolerated at all doses, with a No Observed Adverse Effect Level established at 1.2E11 vg/eye, supporting its clinical advancement for USH1B.

## INTRODUCTION

Usher syndrome type 1B (USH1B) is the most severe form of Usher Syndrome and accounts for more than 50% of all USH1 cases^1^. It is caused by mutations in *MyosinVIIa* (*MYO7A*), which encodes an unconventional myosin protein that functions in hair cells of the inner ear and in the retina^2-5^. Affected patients are born profoundly deaf with vestibular dysfunction and typically present with progressive retinitis pigmentosa (RP) in their first decade of life^6^. Although cochlear hair cells of the inner ear are absent or malformed at birth, cochlear implants have proven successful in restoring hearing to USH1B patients, especially when implanted at a young age^7^. Unfortunately, no such treatment is currently available for the progressive vision loss associated with this disease. Previous studies show that USH1B-associated RP varies in age of onset and severity depending on the underlying pathogenic *MYO7A* mutation. However, in most cases, central and peripheral islands of structurally and functionally normal retina are maintained for years^8-11^. Taken together with the fact that newborn hearing screening leads to early genetic confirmation of *MYO7A*-associated USH1B, these patients may be good candidates for gene replacement therapy.

In 2014, the UshStat trial which evaluated an Equine Infectious Anemia Virus (EIAV)-based lentivirus to deliver full length *MYO7A* following subretinal injection in patients with advanced disease was initiated (NCT01505062). No evidence for improvement was noted which was likely due to inefficient transduction of the target cell-photoreceptors (PRs). Lentiviral vectors do not efficiently transduce PRs in postmitotic retinas^12-15^. Instead, they predominantly transduce the retinal pigment epithelium (RPE). In primate retinas, MYO7A localizes to the calyceal processes of PRs, and these cells are the initial site of disease in USH1B patients^8,9,11,16-18^. Immunoblotting showed that MYO7A is expressed at much higher levels in primate neural retina than RPE^19^. In contrast, mice do not have PR calyceal processes and do not exhibit any obvious photoreceptor phenotypes (i.e. degeneration/dysfunction/impaired protein trafficking through the PR connecting cilium)^16,19^. In mouse retina, MYO7A is expressed predominantly in the RPE, and its absence is associated with impaired melanosome migration in these cells^19,20^. The differential expression of MYO7A between mice and man highlights the lack of suitability for incorporating RPE cell function as a surrogate for clinical efficacy, and the inability of lentiviral vectors to transduce PRs points to their futility for the successful treatment of USH1B.

Adeno-associated virus (AAV) has emerged as the gold standard for targeting therapeutic genes to the retina, and in particular, photoreceptors. However, its relatively small packaging capacity (∼4.8 kb) does not accommodate the *MYO7A* cDNA (∼6.7 kb). We, and others, have overcome this hurdle by developing dual AAV vector platforms that promote expression of large genes via different mechanisms^21^. The DNA-recombination approach involves splitting the large cDNA into front and back half vectors that are co-delivered and recombine via regions of complementary DNA^22-24^. In the simplest, “overlap” dual vector system, front and back half vectors both contain an identical, shared cDNA sequence derived from the gene of interest that drives recombination^22,25^. In the “hybrid” dual vector system (aka Dual AAV-trans splicing), the front half vector contains the 5’ cDNA, a splice donor site and homologous recombination sequence (HRS) and the back half vector contains complementary HRS, a splice acceptor site and the 3’ cDNA. Recombination of hybrid dual vectors occurs via the HRS or AAV ITRs and, after production of mRNA, these sequences are spliced out^22,26^. This approach is being used to address *MYO7A*-associated USH1B in a Phase 1/2 clinical trial (NCT06591793). In the protein trans-splicing approach (aka ‘split-intein’) the cDNA is split across two independent vectors designed to create N’ or C’ terminal polypeptides. Within each peptide is one half of a split-intein which bridges the two polypeptides together after translation and then spontaneously self-excises. This approach is being used by multiple sponsors (NCT07161544 and NCT06942572) in Phase 1/2 trials to address *ABCA4*-associated Stargardt disease (STGD). In the mRNA trans-splicing approach, each vector results in transcription of an independent mRNA (encoding the front and back half of the gene of interest) a complementary binding domain (BD) to promote recombination and splice donor/acceptor sites that ultimately remove BD sequence from the mature mRNA. This approach is also being used to address *ABCA4*-associated STGD (NCT07002398) in a Phase 1/2 trial. Notably, due to their inherent design, both split-intein and RNA trans-splicing approaches generate truncated polypeptides or transcripts, and there is the potential for the production of non-target proteins.

We previously showed that both overlap and hybrid dual AAV vectors drive full length MYO7A expression *in vitro* and in subretinally injected C57BL/6J mice^22^. While hybrid dual vectors produced the highest levels of full-length MYO7A in the mouse retina, they also produced a truncated protein derived from the front half vector and, at a relatively high dose (2e10 vg), led to a reduction in retinal thickness, an unwanted safety signal. When delivered at the same dose, overlap dual vectors did not produce truncated protein or cause loss of retinal structure but expression of full length MYO7A was significantly lower than that produced by the hybrid platform. The results suggested that dual AAV vectors with defined genetic payloads are a viable treatment option for USH1B. However, these first-generation (v1) platforms had potential drawbacks and afforded the opportunity for additional optimization. The goal of this study was to optimize the safety and efficiency of our dual AAV vector platforms, to confirm that they led to efficient MYO7A expression in primate photoreceptors, and generate the supporting information needed to advance a laterally spreading dual AAV.SPR-based gene therapy for USH1B into the clinic.

## RESULTS

### First-generation dual AAV vectors drive full length MYO7A in subretinally injected *Myo7a* KO mice

We previously showed that first generation (“v1”) hybrid and overlap dual vectors drove expression of full length MYO7A in subretinally injected “normal”, C57BL/6J mice^22^. However, that study evaluated a relatively high dose of vector [2.0E10 vg total (1.0E10 vg each of front and back half vector)] on a *Myo7a^+/+^* background and was associated with retinal degeneration. Here, v1 vectors were evaluated at a more clinically relevant dose (when scaled for mouse eye) on a MYO7A null background. *Myo7a* KO mice received uniocular subretinal injections with 5E8 vg per eye (2.5E8 vg each of front and back half vector, respectively, n= 6-8 eyes per treatment) of either hybrid-v1 or overlap-v1 dual vectors containing the ubiquitous smCBA promoter and packaged in AAV8(Y733F) capsid (**Figure 1**).

**Figure 1.**
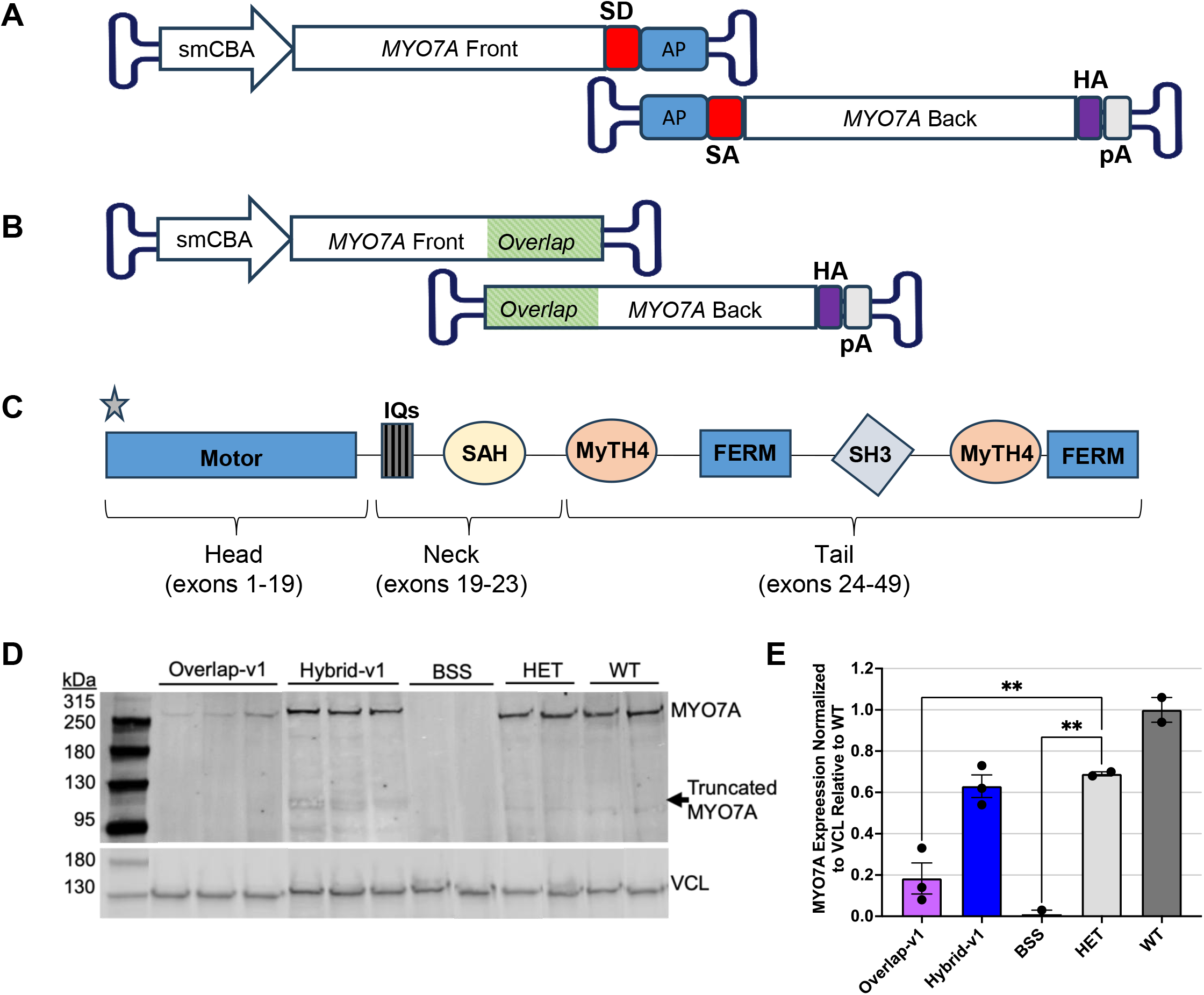
Testing of hybrid-v1 and overlap-v1 dual vectors. (A) Schematic of hybrid-v1 dual vectors. The front half vector contains the promoter, 5’ *MYO7A* cDNA, an intronic element containing a splice donor (SD) sequence, and alkaline phosphatase (AP) sequence. The back half vector contains AP sequence, an intronic element containing a splice acceptor (SA) sequence, 3’ *MYO7A* cDNA, an HA tag, and a bGH polyA. (B) Schematic of overlap-v1 dual vectors. The front half vector contains the promoter and 5’ *MYO7A* cDNA. The back half vector contains 3’ *MYO7A* cDNA, an HA tag, and bGH polyA. Complimentary sequence (1365 bp) shared between both vectors is designated by green boxes. (C) Schematic of MYO7A protein sequence. The head consists of the motor domain and corresponds to exons 1-19 of the gene. The neck is comprised of five IQ sites and a SAH, which corresponds to exons 19-23 of *MYO7A*. The tail contains two MyTH4-FERM regions interspersed by a single SH3 motif and corresponds to exons 24-49 of the gene. The approximate binding site of the anti-MYO7A antibody used in this study (Santa Cruz, sc-74516) is denoted with a star. The antibody binds to amino acids 11-70 within exon 1 and allows for detection of both full length and truncated front-half MYO7A proteins. (D) Western blot showing expression of dual AAV8(Y733F)-mediated MYO7A in subretinally injected *Myo7a* KO mice. Both overlap-v1 and hybrid-v1 dual vectors produced full length MYO7A (∼250 kDa). Truncated MYO7A (∼116 kDa) is detectable only in mice injected with hybrid-v1 dual vectors. (E) Quantification data in D. The hybrid-v1 dual vectors produce full length MYO7A at levels equivalent to that seen in *Myo7a^+/-^*(HET) mice (∼65% of WT expression). Statistical comparisons were performed using a two-way ANOVA with multiple comparisons (*Myo7a^+/-^*HET mice as control group). smCBA= small cytomegalovirus/chicken ß-actin promoter; HA= hemagglutinin tag; pA= bovine growth hormone polyadenylation sequence; SD= splice donor; AP= alkaline phosphatase/AP head sequence; SA= splice acceptor; IQ= calmodulin-binding motif; SAH= single alpha helix; MyTH4-FERM= myosin tail homology 4-band 4.1 ezrin, radixin, moesin; SH3= SRC homology domain 3; kDa= kilodalton; BSS= balanced salt solution/vehicle injection in *Myo7a* KO mice; HET= heterozygous *Myo7a^+/-^*mice; WT= *Myo7a^+/+^* mice; VCL= Vinculin; **p<0.01

At six-weeks post-injection, outer nuclear layer (ONL) thickness in *Myo7a* KO mice, assessed by optical coherence tomography (OCT), was comparable across overlap-v1, hybrid-v1, and BSS-injected groups, with no statistically significant differences observed (**Figure S1**). Full length MYO7A expression (∼250 kDa) was observed in eyes injected with both overlap-v1 and hybrid-v1 dual vectors (**Figure 1D**, n= 2-3 eyes per group). Consistent with our prior findings, hybrid-v1 dual vectors produced significantly more full-length protein than overlap-v1 vectors, at levels equivalent to those seen in heterozygous *Myo7a^+/-^*(HET) mice (**Figure 1E**). Truncated MYO7A (∼116kDa) was detectable only in eyes injected with hybrid-v1 dual vectors, corresponding to the protein expected to be translated from the promoter-containing front half vector.

To further characterize each platform, we evaluated vector genomes and *MYO7A-*derived transcripts corresponding to regions of *MYO7A* delivered in the front or back vectors, respectively, in eyes injected with front half or back half vector alone. When front half vectors were delivered, front half vector genomes and front half vector-derived *MYO7A* transcript were detected, regardless of platform (**Figure S2A, B**). Despite similar levels of front half-derived transcript between platforms, only hybrid-v1 front half vectors produced detectable truncated protein by western blot (**Figure S2E**). In back half vector treated eyes, back half vector genomes were detectable in all eyes, but the levels of back half vector-derived transcript were negligible, a finding consistent with the back half vector lacking a promoter sequence to drive transcription (**Figure S2C, D, E**).

Taken together, these results confirmed the utility of both dual vector platforms for reconstituting full length MYO7A expression but identified key drawbacks requiring further optimization: specifically, the need to (1) reduce truncated protein production from the hybrid platform front half vector, and (2) increase the efficiency of the overlap platform.

### Optimized hybrid dual vectors produce less truncated MYO7A and higher levels of full length MYO7A *in vitro*

Multiple modifications were made to the hybrid-v1 dual vector constructs with the aim of (1) reducing the likelihood that truncated protein produced from the front half vector would have a toxic gain of function, (2) reducing or eliminating translation of truncated protein from the front half vector, and (3) improving packaging efficiency. A summary of these modifications is provided in **Table S1**.

First, we altered the cDNA split point between the front and back half vectors. In hybrid-v1 vectors, the cDNA was split between exons 23/24, corresponding end of the neck and start of tail domain. In our optimized hybrid “v2" vectors, the split point was moved upstream to between exons 21/22. This modification was designed to prevent production of a truncated MYO7A containing the full neck domain and any portions of the tail domain by the front half vector, alone. Structural models and functional analysis of MYO7A and related myosins indicate that the neck and tail domain are required for proper localization and cargo binding^27^. As all of the tail and a portion of the neck domain are absent from the revised exon 21/22 split point front vector, any potential product is predicted to be incapable of localizing to the connecting cilium or competing with endogenous MYO7A for its binding partners, reducing the likelihood of dominant-negative activity. This is supported by comparisons of USH1B patient phenotypes, where those patients with truncation mutations that result in only the motor domain being intact lead to a less severe disease phenotype compared to point mutations in the neck and tail domain^10^. Furthermore, DFNA11, the only autosomal dominant *MYO7A*-associated condition that impairs hearing is characterized by mutations in the tail domain^28,29^. We also removed excess non-coding sequence, and optimized linker and polyA sequences to better fit the transgene elements within the conventional size of rAAV genome. This generated hybrid candidate “v2a," “v2b," “v2c" front variants and hybrid “v2b", “v2 MIN back" and “v2b MIN back” constructs. All back half vectors were generated with or without a C-terminal HA tag, the latter allowing for detection of MYO7A-HA on a normal (e.g. *Myo7a^+/+^*) mouse or non-human primate background.

All optimized hybrid vector combinations were packaged in AAV2 and evaluated in HEK293 cells by co-transduction. All hybrid vector pairs produced full length MYO7A (n= 3-11 replicates per treatment) (**Figure 2**). Several optimized combinations produced significantly higher levels of full-length protein relative to hybrid “v1” vectors, including "v2 + v2 MIN" (p<0.0001) and "v2c + v2b MIN" (p<0.01) (**Table S2**). All optimized hybrid vector pairs produced less truncated MYO7A relative to first generation vectors, with the truncated product running at a slightly smaller molecular weight than that produced by hybrid “v1” vectors, consistent with the upstream shift in the cDNA split point (**Figure 2A, C**). These data support the notion that (1) additional sequence-level optimization of the front half vector further reduces truncated protein production and (2) reducing the size of the back half vector construct increases the efficiency of the hybrid dual vector platform.

**Figure 2.**
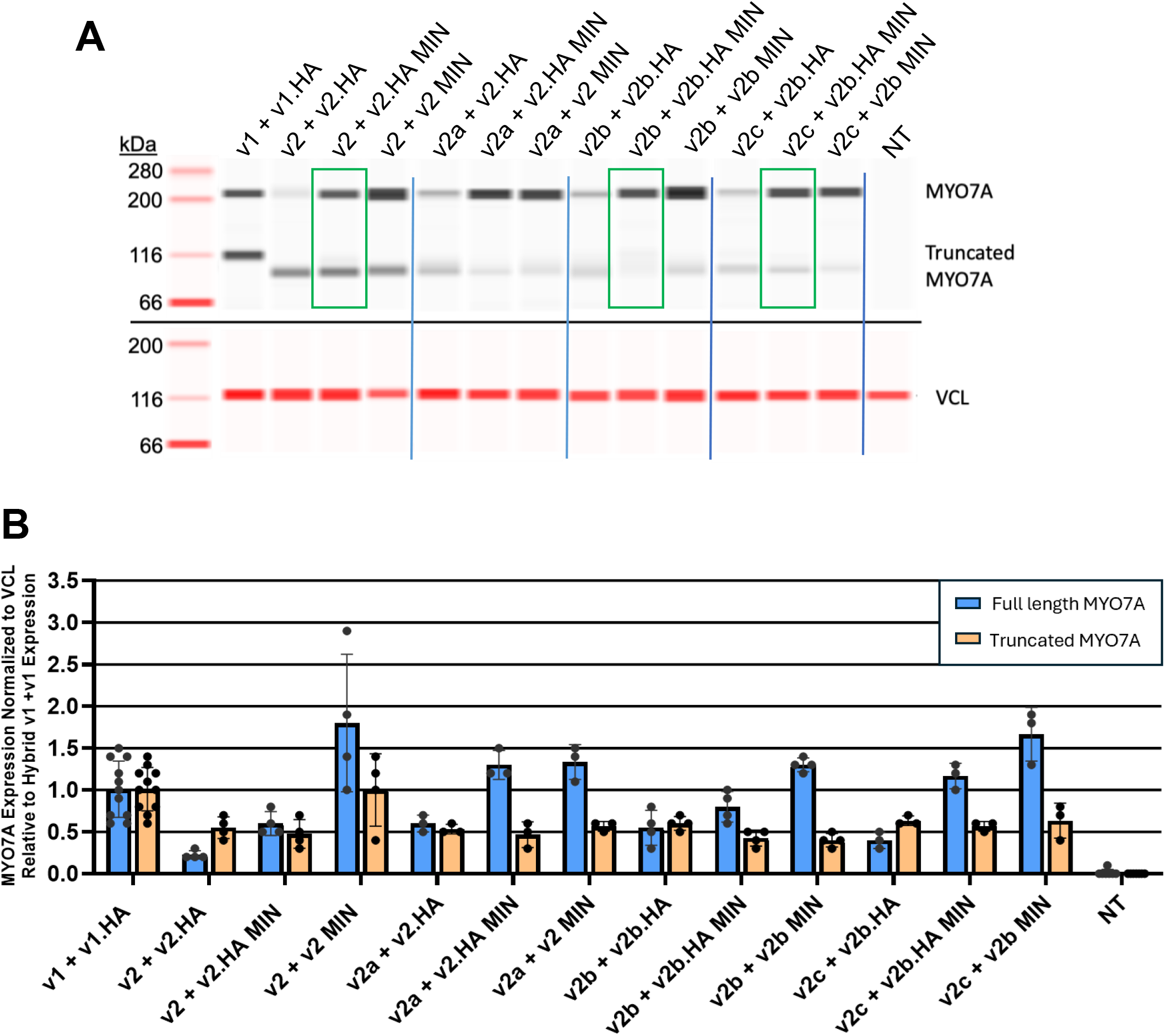
*In vitro* testing of optimized hybrid dual vector sets. (A) Representative Jess western blot image of HEK293 cells co-transduced with different hybrid dual vector sets (n = 3–11 replicates per treatment; the highest-expressing samples from each dual vector pair are shown). All vectors were packaged in AAV2. Vector pairs are labeled above each lane according to their front and back half vector combination. Green boxes highlight the lead vector pairs selected for further evaluation based on their combination of robust full-length MYO7A expression and reduced truncated protein production. Full-length MYO7A (∼225 kDa) and truncated front-half MYO7A are detected in the upper panel; note that truncated protein produced by optimized vectors runs at a smaller molecular weight than that produced by hybrid v1 + v1 vectors. Vinculin (VCL, ∼116 kDa) is shown as a loading control in the lower panel. (B) Quantification of full-length and truncated MYO7A expression resulting from each dual vector pair, normalized to VCL and expressed relative to hybrid v1 + v1 expression. Several optimized hybrid vector combinations produced significantly higher levels of full-length MYO7A protein relative to first-generation hybrid v1 + v1 vectors, including v2 + v2 MIN (p<0.0001) and v2c + v2b MIN (p<0.01). All but one optimized hybrid vector pair produced less truncated MYO7A relative to first-generation vectors. Statistical comparisons were performed using a two-way ANOVA with multiple comparisons (hybrid v1 + v1 as control group). kDa = kilodaltons; VCL = Vinculin; *p<0.05; **p<0.01; ****p<0.0001.

### Optimized overlap dual vectors drive higher levels of full-length MYO7A *in vitro*

Our first-generation overlap-v1 front and back half vectors contain 1,365 bp of overlapping *MYO7A* cDNA sequence. We hypothesized that reducing this overlap length might improve recombination efficiency and thereby increase full length MYO7A expression. Toward that end, we designed additional vector constructs that, when paired, contained successively smaller overlap lengths spanning a broad range (**Table S3**). All back half vectors were generated with a C-terminal HA tag. All optimized overlap constructs were packaged in AAV2 and co-transduced in HEK293 cells (n= 4–6 replicates per treatment).

All but one overlap vector pair produced full length MYO7A (**Figure 3**). Progressively reducing the overlap length resulted in increasing levels of full length MYO7A expression. The overlap “v3+v3” combination which contained an intermediately sized overlap length produced the highest levels of full-length protein, reaching levels equivalent to those achieved by hybrid “v1” vectors (p<0.0001 vs. overlap v1 + v1) (**Figure 3B**). Further reduction of the overlap length below this point caused full-length MYO7A expression to drop back to levels seen with the original v1 overlap vectors, and truncated protein became detectable (**Figure 3A**). The overlap “v2+v3” pair, which had the shortest overlap length tested, produced no full-length protein. These data demonstrate that reducing the length of overlapping *MYO7A* cDNA sequence can increase the efficiency of overlap dual vectors, but only within an optimal range; excessive reduction in overlap length is detrimental to recombination efficiency and introduces truncated protein production.

**Figure 3.**
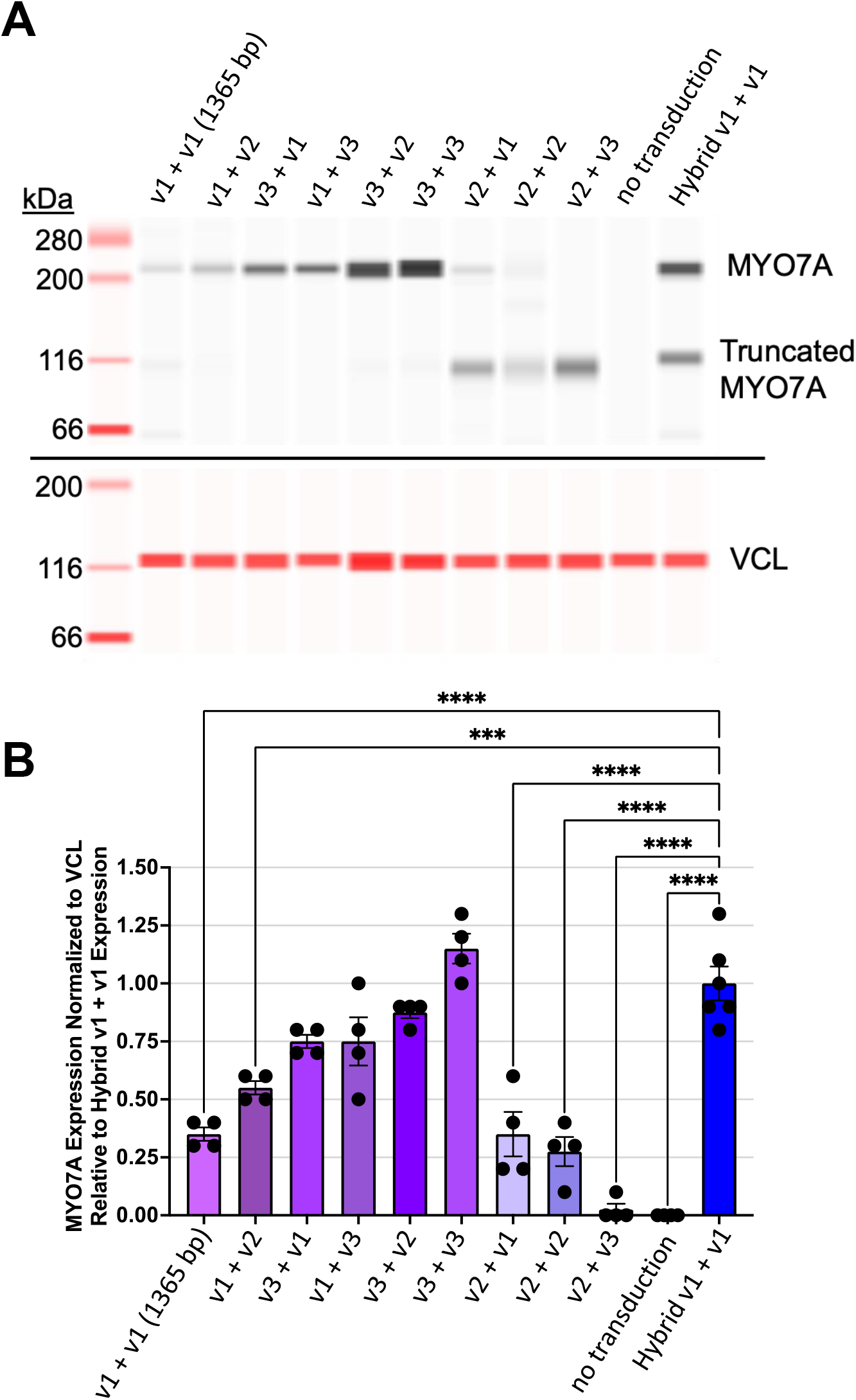
*In vitro* testing of optimized overlap dual vector sets. (A) Representative Jess western blot image of HEK293 cells co-transduced with different overlap dual vector pairs (n = 4–6 replicates per treatment; the highest-expressing samples from each dual vector pair are shown). All vectors were packaged in AAV2. Vector pairs are labeled above each lane according to their front and back half vector combination, with the length of overlapping MYO7A cDNA sequence (bp) indicated in parentheses. Overlap lengths tested were progressively reduced from the original 1,365 bp (v1 + v1) to the shortest sequence (v2 + v3). The first-generation hybrid v1 + v1 vector pair is included as a reference comparator in the final lane. Full-length MYO7A (∼225 kDa) is detected in the upper panel; vinculin (VCL, ∼116 kDa) is shown as a loading control in the lower panel. Truncated MYO7A (∼116 kDa) is detectable in overlap vector pairs with the short overlap lengths (v2 + v1; v2 + v2), and no full-length MYO7A is produced by the v2 + v3 pair. (B) Quantification of full-length MYO7A expression resulting from each overlap dual vector pair, normalized to VCL and expressed relative to hybrid v1 + v1 expression. Progressively reducing the overlap length from 1,365 bp resulted in increasing levels of full-length MYO7A expression, with the v3 + v3 combination producing the highest levels of full-length protein, reaching levels equivalent to those achieved by hybrid v1 + v1 vectors. Further reduction of overlap length resulted in decreased full-length MYO7A expression, with truncated protein becoming detectable. Statistical comparisons were performed using a two-way ANOVA with multiple comparisons (hybrid v1 + v1 as control group). kDa = kilodaltons; VCL = Vinculin; ***p<0.001; ****p<0.0001.

### Optimized dual vector constructs are well tolerated and drive robust full-length MYO7A expression in subretinally injected *Myo7a* KO mice

Based on the *in vitro* optimization data, hybrid vector pairs ("v2 + v2.HA MIN," "v2b + v2b.HA MIN," and "v2c + v2b.HA MIN") and overlap pairs ("v3 + v2" and "v3 + v3") were advanced to *in vivo* testing in *Myo7a* KO mice. All constructs contained HA tags that would ultimately allow for staining of vector-derived MYO7A-HA in non-human primate retinas that express endogenous MYO7A. Vectors were packaged in the recently characterized AAV.SPR capsid for its superior performance, including in primate retina^30^. The first generation “v1 + v1” hybrid vector pair was packaged in AAV8(Y733F), to serve as a basis of comparison to prior experiments. Vectors were administered at a 1:1 ratio (5E8 vg each, 1.0E9 vg total) via subretinal injection (n= 7-11 eyes per treatment). Animals were evaluated at six weeks post-injection.

All hybrid and overlap vector pairs produced robust full length MYO7A expression in injected eyes (**Figure 4A**). All three optimized hybrid pairs demonstrated significantly reduced truncated MYO7A expression compared to the first-generation hybrid “v1+v1” (p<0.001) (**Figure 4A, C**). OCT analysis revealed no significant differences in ONL thickness across any treatment group relative to BSS controls, confirming the absence of vector-associated retinal toxicity at this dose (**Figure 4D**). Immunohistochemical analysis using an anti-HA antibody revealed MYO7A-HA signal localized to the photoreceptor inner segment/outer segments (IS/OS) and retinal pigment epithelium (RPE) across all treatment groups (**Figure S3**).

**Figure 4.**
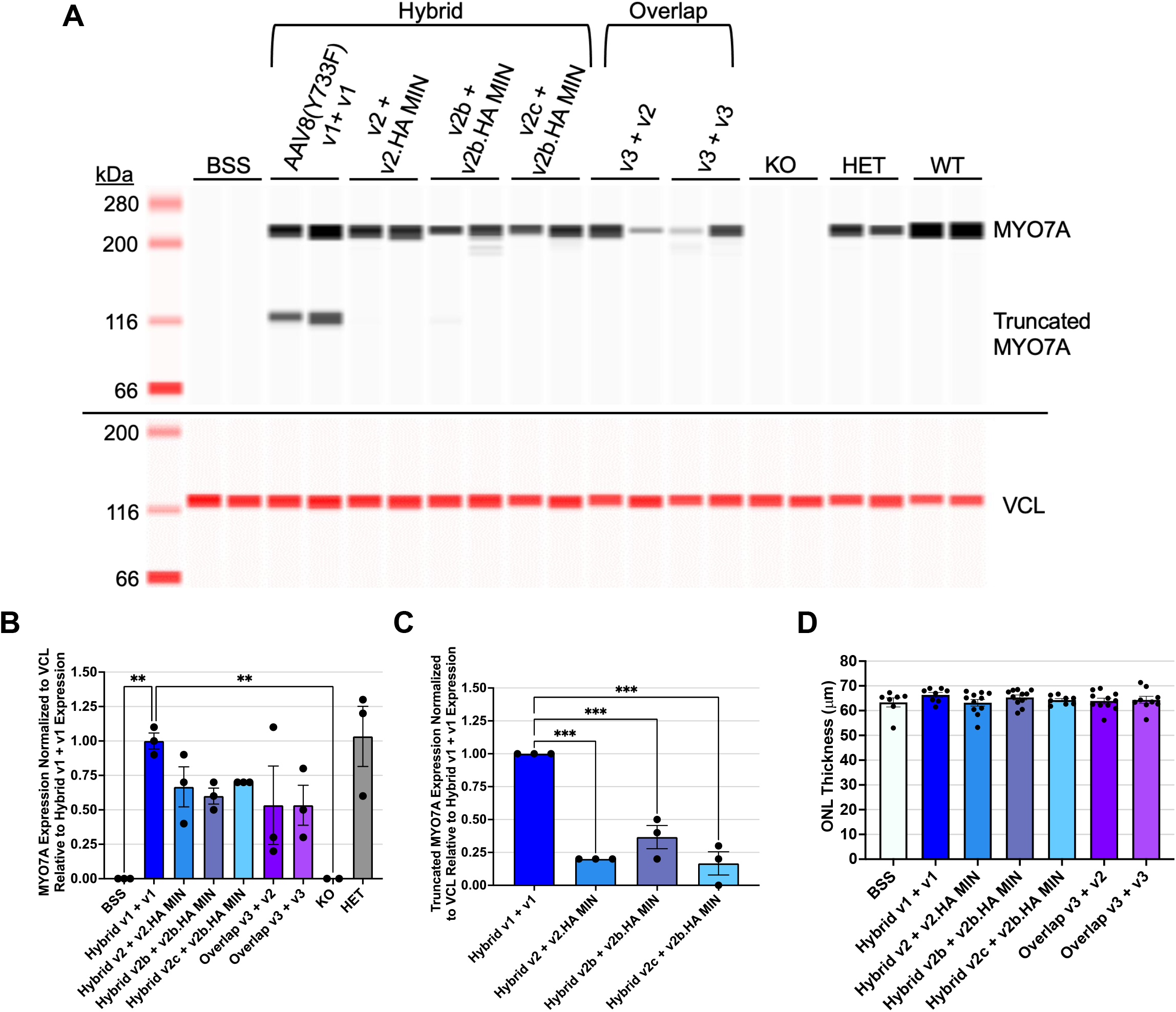
*In vivo* testing of optimized hybrid and overlap dual vector pairs in subretinally injected *Myo7a* KO mice. (A) Representative Jess western blot image of retinal lysates from *Myo7a* KO mice probed with an antibody directed against MYO7A (n = 4–8 eyes per treatment group; the two highest-expressing samples from each dual vector pair are shown). All vectors were packaged in AAV.SPR except for the first-generation hybrid v1 + v1 vectors, which were packaged in AAV8(Y733F). BSS-injected *Myo7a* KO (KO), heterozygous *Myo7a^+/-^*(HET), and *Myo7a^+/+^* mice (WT) mice served as controls. Full-length MYO7A (∼225 kDa) and truncated front-half MYO7A (∼116 kDa) are detected in the upper panel; vinculin (VCL, ∼116 kDa) is shown as a loading control in the lower panel. (B) Quantification of full-length MYO7A expression resulting from each dual vector pair, normalized to VCL and expressed relative to hybrid v1 + v1 expression. All hybrid and overlap vector pairs produced full-length MYO7A expression. (C) Quantification of truncated front-half MYO7A expression resulting from each hybrid dual vector pair, normalized to VCL and expressed relative to hybrid v1 + v1 truncated expression. All three optimized hybrid pairs demonstrated significantly reduced truncated MYO7A expression compared to the first-generation hybrid v1 + v1. (D) Average outer nuclear layer (ONL) thickness (µm) measured by optical coherence tomography (OCT) across all treatment groups. No significant differences in ONL thickness were observed relative to BSS-injected controls, confirming the absence of vector-associated retinal toxicity at this dose. Statistical comparisons in panels B and C were performed using a two-way ANOVA with multiple comparisons (hybrid v1 + v1 as control group). Statistical comparisons in panel D were performed using a two-way ANOVA with multiple comparisons (BSS as control group); no significant differences were identified. kDa = kilodaltons; VCL = Vinculin; BSS = balanced salt solution; KO = *Myo7a^-/-^*homozygous knockout mice; HET = heterozygous *Myo7a^+/-^*mice; WT = *Myo7a^+/+^* mice; **p<0.01; ***p<0.001.

Taken together, these results confirmed that multiple optimized hybrid and overlap dual vector pairs drive robust, safe MYO7A expression in the *Myo7a* KO mouse retina. The hybrid “v2c + v2b.HA MIN” and overlap “v3 + v3” pairs were selected as the top candidates for subsequent capsid comparison studies.

### AAV.SPR outperforms AAV8-based dual MYO7A vectors in subretinally injected *Myo7a* KO mice

Next, we evaluated the relative efficiency of the optimized hybrid (“v2c + v2b.HA MIN”) and overlap (“v3 + v3”) vector pairs in context of AAV8(Y733F) vs. AAV.SPR following subretinal injection in *Myo7a* KO mice. For all subsequent studies, "Opt Hybrid" refers to v2c + v2b.HA MIN and "Opt Overlap" refers to v3+v3. Front and back half vectors were delivered at 1:1 ratio (5E8 vg each, 1.0E9 vg total).

AAV.SPR Opt Hybrid vectors produced the highest levels of full length MYO7A among all groups tested, reaching ∼1.4-fold greater expression than HET controls (p<0.001 vs. HET) (**Figure 5A, B**). While truncated MYO7A was detectable across Opt Hybrid-injected groups, levels expressed as a percentage of full-length protein were low (an average of ≤15%) and did not differ between AAV.SPR and AAV8(Y733F) hybrid vectors (**Figure 5C**). Immunohistochemical analysis confirmed robust MYO7A-HA signal in photoreceptor IS/OS and RPE in AAV.SPR Opt Hybrid-injected eyes, with signal intensity visibly greater than that observed in AAV8(Y733F) Opt Hybrid, AAV.SPR Opt Overlap, or AAV8(Y733F) Opt Overlap groups (**Fig S4**). These results further supported the selection of AAV.SPR as the preferred capsid for the clinical candidate.

**Figure 5.**
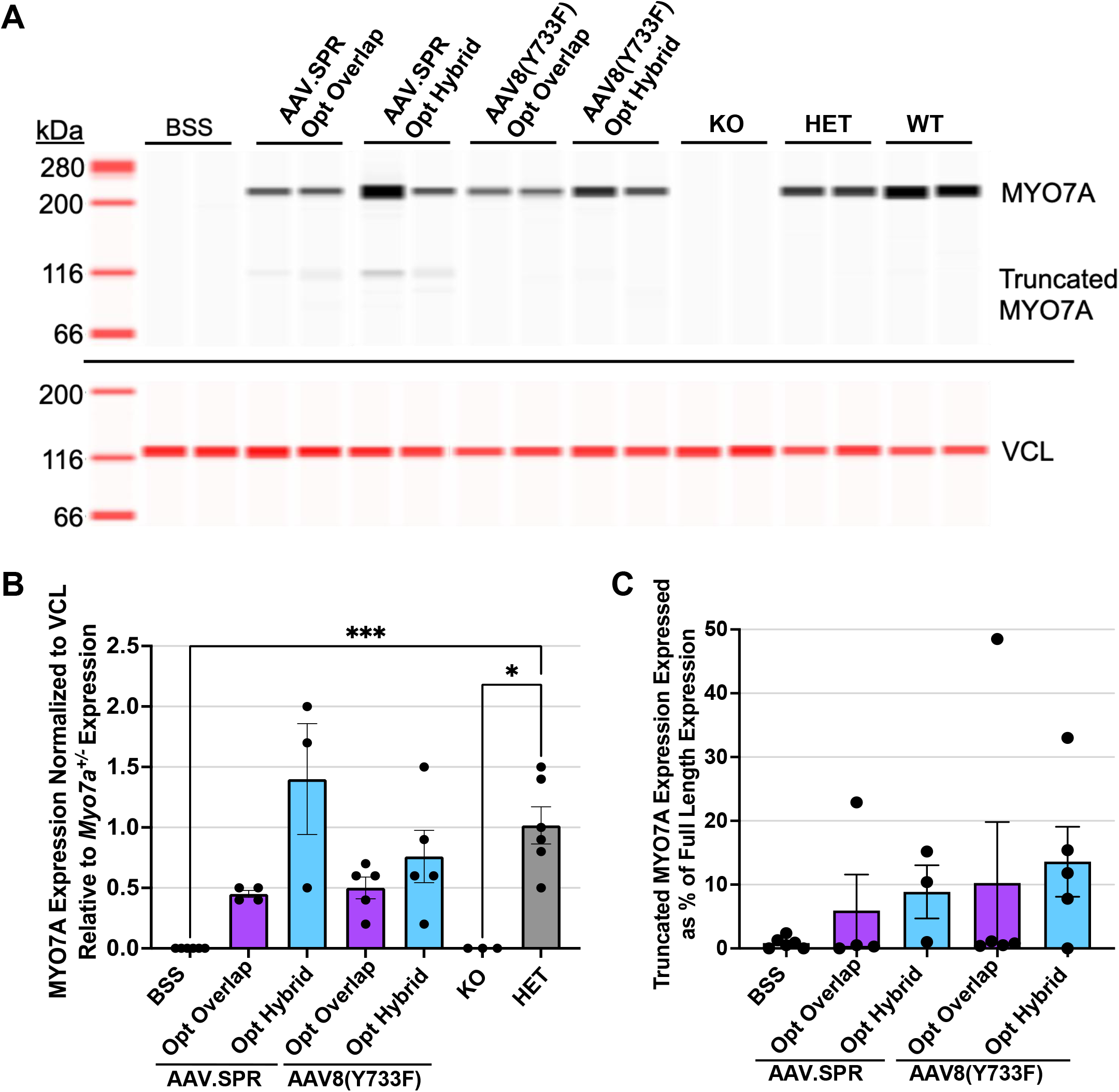
AAV.SPR outperforms AAV8(Y733F)-based dual vectors in subretinally injected Myo7a KO mice. (A) Representative Jess western blot image of retinal lysates from *Myo7a* KO mice subretinally injected with the optimized hybrid (v2c + v2b.HA MIN, aka Opt Hybrid) or overlap (v3 + v3, aka Opt Overlap) dual vector pairs packaged in either AAV.SPR or AAV8(Y733F). BSS-injected *Myo7a* KO (KO), heterozygous *Myo7a^+/-^*(HET), and *Myo7a^+/+^* (WT) mice served as controls (n=3-6 eyes). Full-length MYO7A (∼225 kDa) and truncated front-half MYO7A (∼116 kDa) are detected in the upper panel; vinculin (VCL, ∼116 kDa) is shown as a loading control in the lower panel. (B) Quantification of full-length MYO7A expression from each treatment group, normalized to VCL and expressed relative to *Myo7a^+/-^*HET expression. (C) Truncated MYO7A expression expressed as a percentage of full-length MYO7A expression for each treatment group. No statistical comparisons were performed for truncated protein. For all subsequent studies, “hybrid" refers to the v2c + v2b.HA MIN vector pair and “overlap" refers to the v3 + v3 vector pair. Statistical comparisons were performed using a two-way ANOVA with multiple comparisons (HET as control group). kDa = kilodaltons; VCL = Vinculin; BSS = balanced salt solution; KO = *Myo7a^-/-^*homozygous knockout mice; HET = heterozygous *Myo7a^+/-^*mice; WT = *Myo7a^+/+^* mice; *p<0.05; ***p<0.001.

### AAV.SPR dual MYO7A vectors drive supraphysiological levels of MYO7A in non-human primate retina

Next, we evaluated the dose response of AAV.SPR Opt Hybrid and AAV.SPR Opt Overlap dual vectors in cynomolgus monkeys subretinally injected with high (1.0E11 vg/eye), mid (4.0E10 vg/eye L), or low doses (1.6E10 vg/eye) of each vector pair (**Table S4**). Two distinct 75 μL subretinal blebs were created per eye, placed peripherally to avoid foveal detachment. Both Opt Hybrid and Opt Overlap dual vectors were well tolerated following subretinal injection. Mild reactions to the subretinal surgery including RPE pigment alteration and partial loss of the ellipsoid zone detail on OCT were common across all groups, including vehicle controls and were consistent with what has been observed following subretinal administration^31^. Uveitis scores were similar between platforms and were mild and transient. **Table S5** lists only the days when findings were noted. All other timepoints had no findings. No findings were reported in H&E-stained retinal sections of any animal.

Retinal punches were collected from bleb 1, bleb 2, the macula, and a distal region nine weeks post-injection and analyzed by western blot using anti-HA and anti-MYO7A antibodies. HA-tagged MYO7A was detected in retinal punches from animals receiving all doses of AAV.SPR Opt Hybrid vectors, with expression at bleb, macular, and distal sites at the high and mid doses (**Figure 6A, B**). MYO7A-HA signal at the low dose was markedly reduced, with expression primarily restricted to bleb sites. AAV.SPR Opt Overlap vectors showed a similar dose-dependent pattern, with MYO7A-HA expression levels comparable to AAV.SPR Opt Hybrid across matched dose groups. MYO7A protein levels in treated NHP eyes were elevated 2 to 10-fold above vehicle-injected controls across all dose groups for both platforms, with similarly high expression observed at the high doses of Opt Hybrid and Opt Overlap vectors (**Figure 6C, D**). Immunohistochemical analysis confirmed HA-tagged MYO7A expression in peripheral and macular retina (photoreceptors and RPE) in Opt Hybrid and Opt Overlap mid and high dose groups, confirming transduction at clinically relevant doses (**Fig S5**). Although MYO7A expression levels were comparable between platforms in NHP, the consistently superior full-length expression observed across multiple mouse studies supported selection of the AAV.SPR Opt Hybrid dual vectors as the clinical candidate, with a dosing range informed by the safety profile and MYO7A expression levels observed in NHP.

**Figure 6.**
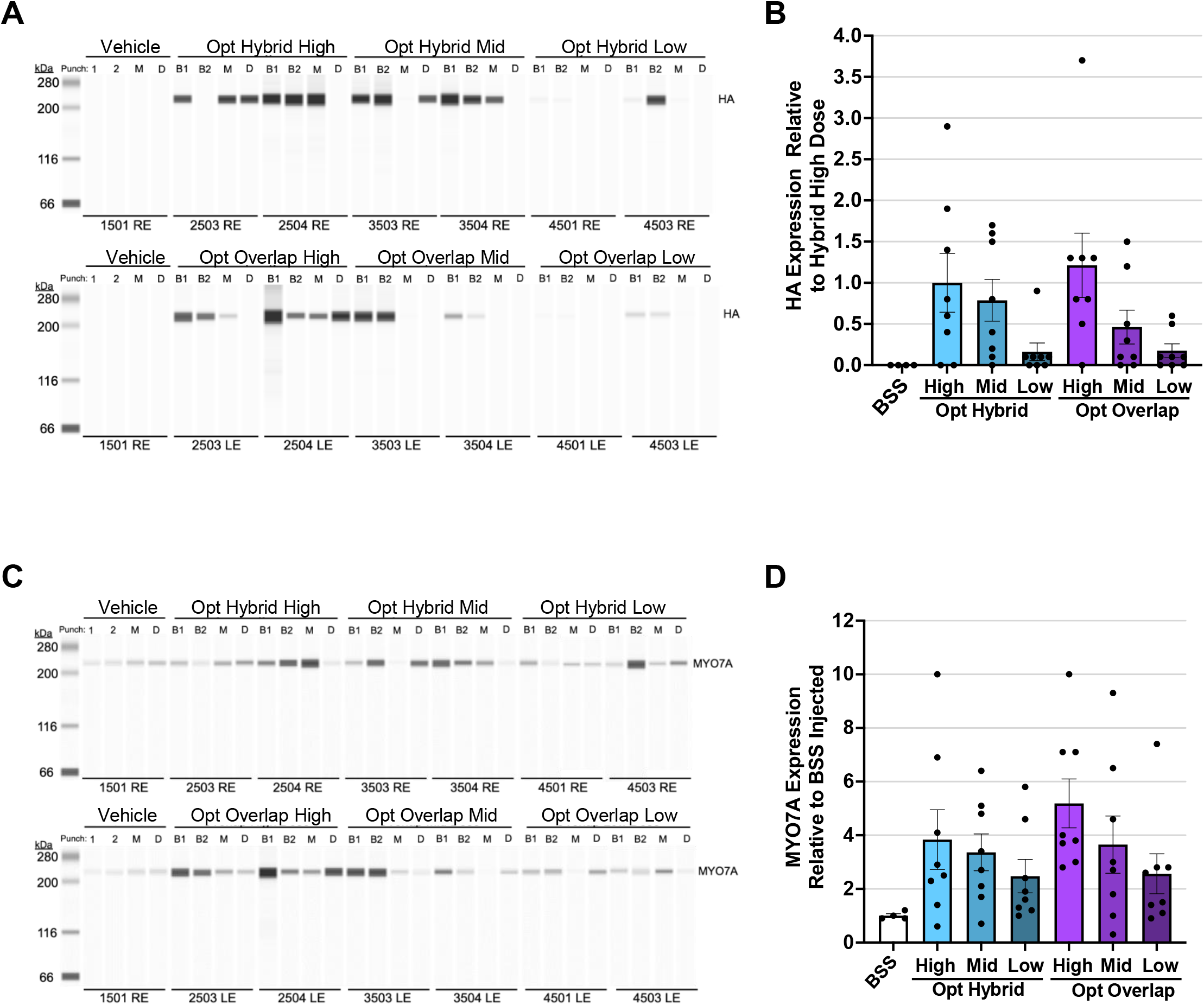
Dose-dependent expression of MYO7A protein following subretinal injection of AAV.SPR Opt Hybrid and Opt Overlap dual vectors in cynomolgus monkeys. (A) Jess western blot images of retinal punches collected from cynomolgus monkeys subretinally injected with AAV.SPR Opt Hybrid (right eye) or AAV.SPR Opt Overlap (left eye) dual vectors at high (6.6E11 vg/mL; 1.0E11 vg/eye), mid (2.64E11 vg/mL; 4.0E10 vg/eye), or low (1.06E11 vg/mL; 1.6E10 vg/eye) doses, probed with an anti-HA antibody. Retinal punches were collected from bleb 1 (B1), bleb 2 (B2), the macula (M), and a distal region (D) of each eye nine weeks post-injection. Vehicle-injected animal (1501 RE) is shown as a negative control. HA-tagged full-length MYO7A (∼225 kDa) is detected in all eyes. HA expression was detected at bleb, macular, and distal sites in hybrid-injected eyes at the high and mid doses, with markedly reduced signal at the low dose. AAV.SPR Opt Overlap vectors showed a similar dose-dependent pattern, with generally lower HA expression levels than AAV.SPR Opt Hybrid across matched dose groups. (B) Quantification of HA-tagged MYO7A expression across all dose groups and retinal punch locations, normalized to total protein and expressed relative to the hybrid high dose group. (C) Representative Jess western blot images of retinal punches from the same animals as in panel A, probed with an anti-MYO7A antibody to detect both vector-delivered and endogenous MYO7A protein (∼225 kDa). (D) Quantification of total MYO7A expression across all dose groups and retinal punch locations, normalized to total protein and expressed relative to vehicle-injected (BSS) controls. MYO7A protein levels in treated eyes were elevated 2– 10-fold above vehicle-injected controls across all dose groups for both platforms. Two blebs of 75 µL each were administered per eye, placed extrafoveally to avoid foveal detachment. Animal numbers are indicated below each bleb group. BSS = balanced salt solution; B1 = bleb 1; B2 = bleb 2; M = macula; D = distal; RE = right eye; LE = left eye; kDa = kilodaltons.

### The safety and biodistribution of ATSN-301 is confirmed in a GLP toxicology study in NHP

A 6-month GLP toxicology study was conducted in cynomolgus macaques to evaluate the safety and biodistribution profile of ATSN-301 (AAV.SPR capsid, smCBA promoter, Opt Hybrid MYO7A dual vectors). Front and back half vectors used in this study were similar to those used in the above non-GLP study barring several minor differences, including use of a kanamycin resistant vector plasmid backbone (as opposed to ampicillin resistant) and removal of the HA tag from the MYO7A cDNA. Animals received bilateral subretinal injections and were assigned to vehicle (n=2), low dose (3.0E10 vg/eye, n=3), mid dose (6.0E10 vg/eye, n=3), high dose (1.2E11 vg/eye, n=3), or high dose unilateral (1.2E11 vg/eye, n=2) groups. An additional non-GLP cohort receiving a version of ATSN-301 containing back vector with HA-tagged MYO7A at the mid dose (n=2) was included per the request of regulators to confirm localization of dual vector-mediated MYO7A (**Table 1**). Two, 75 μL injection blebs were placed extrafoveally to avoid foveal detachment, for a total of 150 μL per eye. Outcome measures included ophthalmic evaluations, intraocular pressure (IOP), electroretinography (ERG), optical coherence tomography (OCT), fundus imaging, clinical pathology, immunogenicity assessment, histopathology, and biodistribution and gene expression analysis of both target and non-target tissues.

**Table 1.**
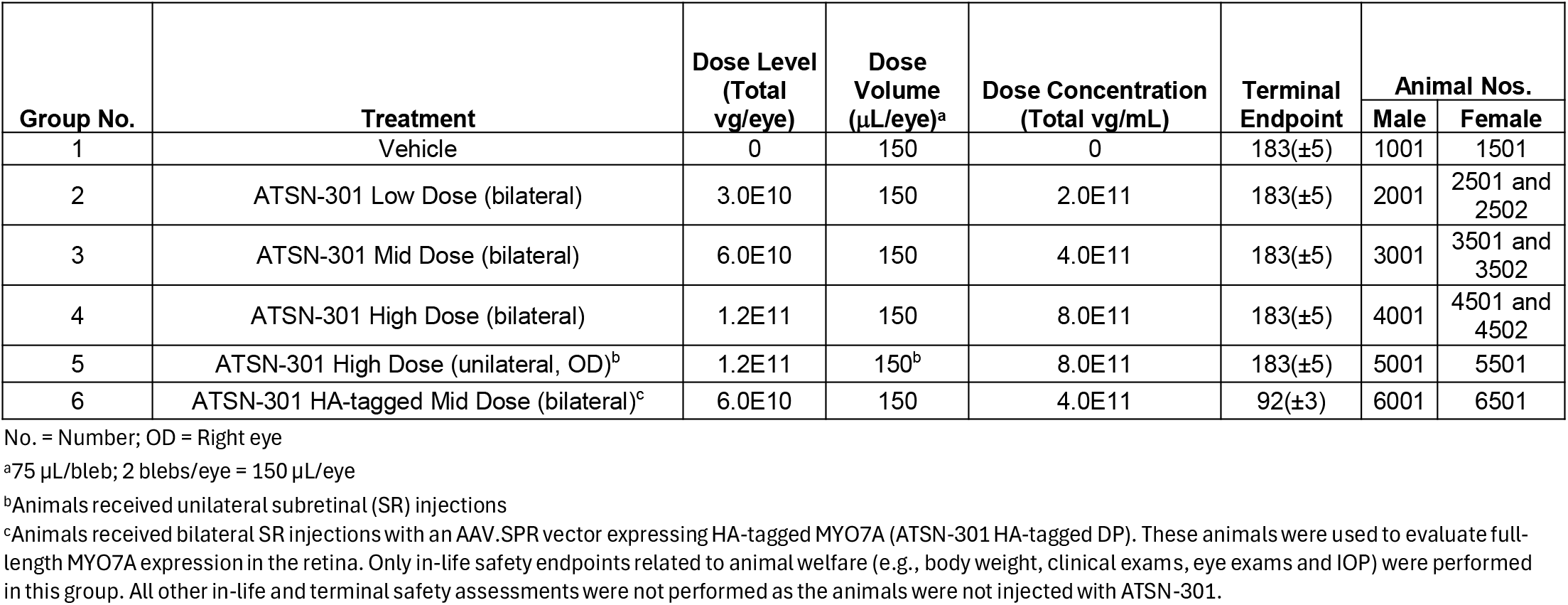

ATSN-301 was well tolerated across all doses evaluated with no treatment-related systemic toxicity. No ATSN-301-related changes were observed in clinical observations, body weight, clinical pathology, or macroscopic findings. There were no scoreable uveitis findings in the vehicle or low-dose groups and no instances of aqueous flare, vitreous cell, vitreous haze, or aqueous/vitreous cell color change at any dose level. Only minimal, low-grade findings were observed in 2 of 11 animals (**Table S6**), consistent with a favorable ocular safety profile for ATSN-301 across the dose range tested. IOP remained within normal limits across all groups and timepoints, with no dose-dependent trends observed. ERG recordings revealed no statistically significant test article-related changes in retinal function at any dose. Ocular findings were limited to dose-dependent minimal to moderate RPE pigment alterations and minimal to mild irregularity or thickening of the RPE layer at higher doses. These observations were corroborated by OCT and histopathology but were confined to the bleb region and were not considered adverse. No treatment-related microscopic findings were identified outside the bleb region. The No Observed Adverse Effect Level (NOAEL) was therefore established at the highest dose tested, 1.2E11 vg/eye.

Anti-drug antibody (ADA) analysis against MYO7A revealed no treatment-related humoral immune responses across all dose groups. The few positive signals observed (e.g., vehicle animals 1001 and 1501, and low dose animal 2502) were transient, low-titer, and not considered clinically meaningful given their occurrence in vehicle-treated controls and lack of dose-dependence. No immunogenicity signal was detected in the mid or high dose groups at any timepoint through the terminal sacrifice.

Biodistribution analysis demonstrated dose-dependent persistence of ATSN-301 vector genomes in retinal tissues, with both front and back vector components detected at levels proportional to administered dose (**Table S7** and **Table S8**). Correspondingly, vector-derived MYO7A mRNA was also detected at high levels in injected eyes across treatment groups, closely tracking the vector genome findings and confirming that the persisting vector genomes were transcriptionally active; mRNA copy numbers in the retina rose with dose, reaching several hundred thousand copies per 100 ng total RNA in high-dose animals, while no mRNA was detected in the contralateral, undosed eyes of the unilateral group (**Table S9**). Beyond the injected retina, mRNA expression was essentially undetectable, with a single low-level exception in the optic nerve of one mid-dose animal; no transcript was found in any other ocular, neural, or systemic tissue, indicating that vector-derived expression remained tightly localized to the retina. Biodistribution beyond the eye was similarly minimal at the vector genome DNA level: pooled anterior segment tissues (ciliary body, iris, and cornea) contained fewer than 1,000 copies; optic nerve samples were below the limit of detection or quantification in all but one animal; and brain regions including the optic chiasm, lateral geniculate nucleus, cerebellum, optic tract, and visual cortex were uniformly below the limits of detection or quantification. In systemic tissues, vector genomes were below the limit of detection or quantification in the great majority of samples, with very low quantifiable levels found in only three samples (one spleen and two submandibular lymph node specimens). Quantifiable levels were detected in vitreous humor from six animals at terminal sacrifice. Blood samples were below the limit of detection or quantification at every timepoint analyzed. These results mirrored the biodistribution pattern of another AAV.SPR based gene therapy vector, ATSN-201, as previously described^32^.

Full-length HA-tagged MYO7A protein was detected by immunoblot analysis in retinal tissue from animals in the non-GLP HA-tagged cohort injected with the mid dose (**Figure 7**). Immunostaining confirmed that MYO7A-HA localized to photoreceptor inner and outer segments and RPE, consistent with the proposed sites of MYO7A function, and expression was detected both within the bleb region and extending into the macula. Minimal truncated MYO7A protein was detected in treated eyes (≤8% of endogenous, full length MYO7A), consistent with the optimized hybrid vector design. The lack of safety signals observed for at least 6 months post-injection supports the notion that this low level of truncated protein did not negatively impact retinal structure or function in these animals.

**Figure 7.**
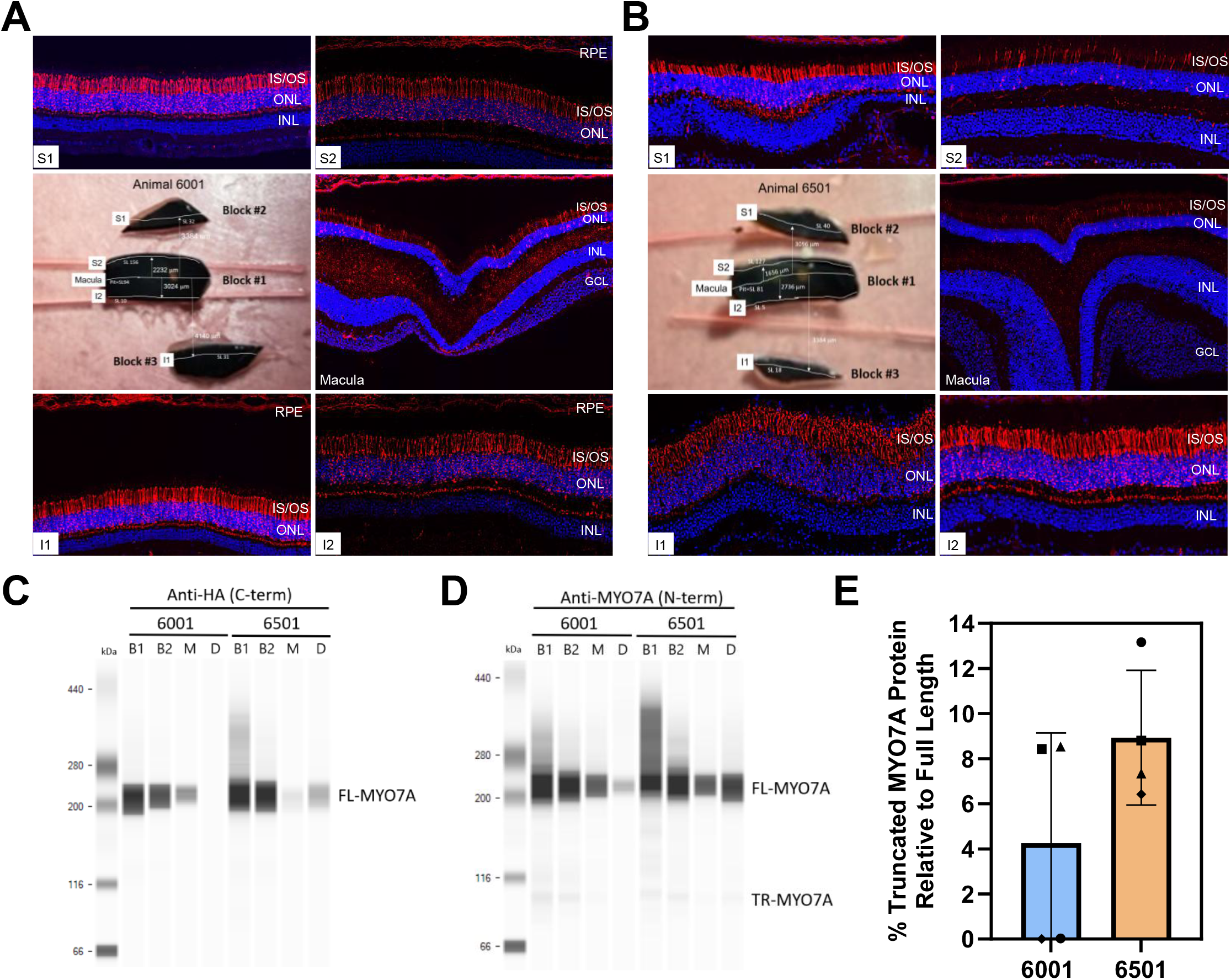
Full-length HA-tagged MYO7A expression in retinal tissue from cynomolgus macaques in the non-GLP HA-tagged cohort of the GLP toxicology study. Animals 6001 and 6501 received bilateral subretinal injections of HA-tagged ATSN-301 at the mid dose (6.0E10 vg/eye) and were terminated at approximately three months post-injection (day 92 ± 3). (A–B) Representative immunohistochemical images of retinal cryosections from Animals 6001 and 6501 stained with anti-HA antibody (red) to detect vector-mediated MYO7A-HA expression and counterstained with DAPI (blue) to label nuclei. HA-tagged MYO7A signal is localized to the photoreceptor inner and outer segments (IS/OS) and retinal pigment epithelium (RPE), consistent with the proposed sites of MYO7A function in the primate retina. Expression is detected both within the bleb region and extending into the macula. (C) Representative Jess western blot of retinal punch lysates from Animals 6001 and 6501 probed with an anti-HA antibody to detect vector-mediated full-length HA-tagged MYO7A (∼225 kDa). Retinal punches were collected from bleb 1 (B1), bleb 2 (B2), the macula (M), and a distal region (D) of the right eye of each animal. (D) Representative Jess western blot of the same retinal punch lysates probed with an anti-MYO7A antibody to detect both full-length HA-tagged and endogenous MYO7A (∼225 kDa) and truncated front-half MYO7A (TR-MYO7A). (E) Quantification of truncated MYO7A expressed as a percentage of full-length MYO7A in retinal punches from Animals 6001 and 6501, measured by Jess western blot. Each MYO7A peak was normalized to the vinculin (VCL) loading control. Individual data points represent the following retinal punch locations: circle = bleb 1, square = bleb 2, triangle = macula, diamond = distal. Truncated MYO7A levels were minimal across all punch locations (up to ∼8% of wild-type MYO7A levels), consistent with the optimized hybrid vector design. IS/OS = inner segment/outer segment; ONL = outer nuclear layer; INL = inner nuclear layer; RPE = retinal pigment epithelium; B1 = bleb 1; B2 = bleb 2; M = macula; D = distal; TR-MYO7A = truncated MYO7A; VCL = vinculin; HA = hemagglutinin; kDa = kilodaltons.

## DISCUSSION

USH1B represents one of the most challenging targets in ocular gene therapy. Its causative gene, *MYO7A*, substantially exceeds the packaging capacity of AAV, and until recently the only clinical attempt to address the associated vision loss was the UshStat trial, in which an EIAV-based lentiviral vector expressing full-length *MYO7A* was administered by subretinal injection to patients with advanced disease (NCT01505062). No evidence of biological activity was noted which, in hindsight, is consistent with the inability of lentiviral vectors to efficiently transduce postmitotic photoreceptors, the primary site of MYO7A function in the primate retina^12-15^. The present study addresses the two fundamental requirements for a successful USH1B gene therapy-1) a vector system capable of delivering full-length *MYO7A*, and 2) a capsid with efficient photoreceptor tropism that avoids the need for foveal detachment in patients with residual central vision.

Our systematic optimization of both hybrid and overlap dual AAV platforms yielded constructs that substantially outperform their first-generation antecedents. For the hybrid platform, modifications to the cDNA split point and additional sequence optimization increased full-length MYO7A output and reduced truncated protein production, the two key liabilities of the v1 design. For the overlap platform, we identified an optimal overlap length that markedly improved recombination efficiency, bringing full-length expression to levels previously achievable only with hybrid vectors. Together, these findings establish clear engineering principles for large-gene dual AAV development that are broadly applicable beyond *MYO7A*. Although both platforms were advanced through optimization and *in vivo* testing, the hybrid platform was selected for clinical development on the basis of consistently superior full-length MYO7A expression which reached supraphysiological levels (∼1.4-fold above heterozygous controls) in the *Myo7a* KO mouse. The residual truncated MYO7A protein produced by the optimized hybrid vectors was low (≤15% of full-length MYO7A in *Myo7a* KO mice; ≤8% of full length, endogenous MYO7A in the GLP NHP study) and was not associated with any adverse findings, supporting the conclusion that the expression advantage of the hybrid platform outweighs the risk posed by truncated protein at the low levels produced. The AAV.SPR capsid provided a meaningful advantage at both stages of *in vivo* evaluation. In the *Myo7a* KO mouse, AAV.SPR-packaged hybrid vectors produced higher full-length MYO7A expression than the equivalent AAV8(Y733F)-packaged vectors. In NHP, AAV.SPR enabled supraphysiological MYO7A expression at macular and distal retinal sites from extrafoveal injection, a clinically important feature that is designed to reduce the surgical risk of foveal detachment in patients with preserved central vision.

The six-month GLP toxicology study confirmed that ATSN-301 was well tolerated across all doses evaluated, with a NOAEL established at 1.2E11 vg/eye, the highest dose tested. Ocular findings were limited to RPE changes confined to the bleb region and were not considered adverse. No ERG changes, systemic toxicity, or meaningful immunogenicity were observed. Biodistribution was restricted to the injected retina, with near-complete absence of vector genomes and transcripts from the optic pathway and systemic tissues, a reassuring profile that aligns with the immune privilege of the subretinal space. The low residual truncated MYO7A detected in treated eyes (∼8% of full length, endogenous MYO7A) produced no histopathological or functional consequences, validating the molecular optimization of the hybrid vector design.

Another dual-AAV MYO7A hybrid vector for USH1B, AAVB-081, has advanced into Phase 1/2 clinical investigation (the LUCE trial, NCT06591793). Published preclinical data for AAVB-081 reveal differences from our approach in capsid, construct design, and dose that provide useful context for interpreting the present findings^33^. ATSN-301 uses the AAV.SPR capsid, which spreads laterally beyond the subretinal bleb margin to reach macular and distal retinal sites from an extrafoveal injection^32^. AAV8, the capsid used in AAVB-081, has not been shown to spread laterally in the primate retina. At the construct level, the AAVB-081 hybrid splits at the exon 24/25 junction, within the MYO7A tail domain, whereas ATSN-301 splits at exon 21/22, excluding the entire tail domain and part of the neck domain from the front-half vector’s coding sequence. With respect to dose, our optimized hybrid vector (the direct progenitor to ATSN-301) elicited over 2-fold the endogenous MYO7A expression level in NHP at 1.6E10 vg/eye. In our GLP toxicology study, the highest dose evaluated, 1.2E11 vg/eye, was established as the NOAEL. By comparison, the NHP toxicology study for AAVB-081 evaluated doses from 1.4E12 to 3.8E12 vg/eye, over 10-fold higher than the ATSN-301 NOAEL. The doses used in the AAVB-081 LUCE trial have not been publicly disclosed. Ferla et al.’s AAVB-081 preclinical publication indicated expected clinical doses of 1.4E11 to 1.4E12 vg/eye—roughly 10-fold higher than the range anticipated for ATSN-301’s first-in-human evaluation^33^.

This study lacks experiments assessing AAV-MYO7A-mediated rescue of retinal function or structure in a model system, due to the absence of models that faithfully mimic human MYO7A retinal biology. MYO7A-deficient mice recapitulate the vestibular and auditory deficits seen in patients but do not undergo retinal degeneration^19,34^. This is unsurprising given the markedly different MYO7A expression pattern between mice and primates: mice predominantly express MYO7A in RPE, while primates and humans predominantly express it in photoreceptors^16,19^. Mice also lack calyceal processes—the fingerlike projections of the inner segment that ring the base of the outer segment^16^. In NHP and humans, MYO7A localizes to these calyceal processes, which are hypothesized to be the photoreceptor analog of stereocilia in vestibular and auditory hair cells^16,35^. Consistent with this, photoreceptors are the initial site of pathology in human USH1B retina^10^. USH1B patient-derived retinal organoids have recently been investigated but, unfortunately, show no phenotypic changes relative to controls^36^. Notably, early versions of what became USHSTAT, the EIAV-based lentiviral gene therapy, reliably rescued RPE melanosome localization in Myo7a mutant mice yet failed in clinical trials, underscoring that mouse-model rescue does not reliably predict clinical efficacy^37^.

Ultimately, clinical application will be essential to establish therapeutic benefit. Nonetheless, the combination of optimized vector design, supraphysiological primate expression from extrafoveal injection, and a clean GLP safety package with well-defined margins provides a strong foundation for the clinical advancement of ATSN-301.

## MATERIALS AND METHODS

### Mouse Model

*Myo7a^-/-^*knockout (*Myo7a* KO) mice were generated and characterized as described previously^19^. Briefly, mouse embryonic stem cells, created as part of the International Knockout Mouse Consortium were acquired from the Sanger Institute, and mice were generated at Jackson Laboratories^38^. Colony founders were backcrossed on a C57BL/6J background (Jackson Laboratory strain#: 000664) for at least five generations prior to use in experiments. The *Myo7a* KO colony was maintained by breeding heterozygous females with homozygous males. C57BL/6J (*Myo7a^+/+^*) controls were generated by breeding heterozygous females with heterozygous males. All mice were genotyped prior to use in experiments using the murine-specific primer sets listed in **Table S10**. The *Myo7a* WT primer set produces an amplicon of 360 bp and the *Myo7a* KO primer set produces an amplicon of 178 bp. The *Myo7a* WT allele was amplified using the following conditions: (1) initial denaturation at 95°C for 2 minutes, (2) 34 cycles of 95°C denaturation for 30 seconds, 66°C annealing for 30 seconds, 72°C elongation for 45 seconds, (3) final elongation at 72°C for 5 minutes. The *Myo7a* KO primer set was run using identical PCR settings, with the exception of a 63°C annealing temperature.

### Animal Care and Ethics Statement

All mice were bred and maintained at the University of Florida Health Science Center Animal Care Services Facility under a 12-hour/12-hour light/dark cycle. Food and water were provided *ad libitum*. All experiments were approved by the University of Florida’s Institutional Animal Care and Use Committee and were conducted in accordance with the Association for Research in Vision and Ophthalmology Statement for the Use of Animals in Ophthalmic and Vision Research and within National Institute of Health regulations.

All procedures performed on Cynomolgus monkeys (*Macaca fascicularis*) were approved by the Charles River Laboratories’ IACUC and performed in accordance with the Association for Research in Vision and Ophthalmology’s Statement for the Use of Animals in Ophthalmic and Vision Research.

### Dual AAV Vector Construct Design

The first-generation MYO7A overlap and hybrid dual vector plasmids were described previously^22^. Front half vector sequences contain the ubiquitous small cytomegalovirus/chicken β-actin (smCBA) promoter^39^, and back half vector sequences contain a hemagglutinin (HA) tag and bovine growth hormone (bGH) polyadenylation signal, unless otherwise noted. The hybrid vectors also contain an 81bp splice donor derived from human hemoglobin subunit beta (HBB) (NC_000011.10) and a 49 bp splice acceptor sequence sharing 29bp in common with mouse IGHV1 NC-000078.7. The 270 bp fragment of alkaline phosphatase (AP) that is present in both front and back half hybrid vectors originates from the human AP gene^40^. Novel dual vector plasmids were generated using traditional molecular cloning methods and sequences confirmed using various sequencing technologies. Additional sequence-level modifications were incorporated into specific optimized front and back half vector constructs, as described in the Results; the precise nature of these modifications can be found in the associated patent application disclosed in the declaration of interests section.

### AAV Vector Production

Research grade vectors were packaged using a standard plasmid-based transfection method in adherent Human embryonic kidney (HEK) 293 cells at Powell Gene Therapy Center (PGTC) Vector Core at the University of Florida. Vectors were purified by iodixanol density gradient centrifugation followed by buffer exchange and concentration using standard methods^41^. *MYO7A* dual vectors intended for use *in vitro* or in mouse were titered using a Pico-Green based protocol as previously described^42^. Vectors intended for *in vitro* studies were packaged in AAV2. Vectors used for mouse and NHP ocular injections were packaged in AAV8(Y733F), or AAV.SPR. Vectors used in NHP experiments were produced at a larger scale (10-stack cell factory, 6,360-cm^2^ cell growth area) and purified as above with the addition of ion-exchange chromatography using a HiTrap Q column (GE Healthcare)^41^. Manufacture of ATSN-301 was performed by a Contract Development Manufacturing Organization using triple transfection of suspension-adapted HEK293 cells. Vectors were purified using AAVX affinity chromatography and cesium chloride density gradient ultracentrifugation followed by dialysis and formulation into BSS-Tween (0.014%). The titer of the front and back vectors were determined by transgene specific ddPCR and combined in a 1:1 ratio to create ATSN-301 drug product. Each undiluted vector preparation used in NHP studies was tested for the presence of endotoxin, with all registering <0.5 EU/mL.

### *In vitro* Transductions

All experiments were performed in HEK293 cells (originally obtained from Dr. Nicolas Muzyczka, University of Florida). Cells were maintained in Dulbecco’s Modified Eagle Media (DMEM) containing 10% fetal bovine serum and 50 mg/mL gentamicin and were incubated at 37°C in 5% CO2. Cell morphology and growth rates were closely monitored throughout passages to maintain near-identical conditions between experiments.

Cells were transduced with virus at a multiplicity of infection (MOI) of 1E4 as previously described^22^. Approximately 24 hours post-transduction, virus-containing media was removed and replaced with complete media. Cells were harvested approximately seventy-two hours post-transduction for analysis via immunoblotting.

### Subretinal Injections

Subretinal injections were performed in *Myo7a* KO mice as previously described^43^. Eyes were dilated with 1% atropine at 30 minutes and 15 minutes prior to injection and 2.5% phenylephrine HCL at 5 minutes prior to injection. Mice were anesthetized with ketamine (100 mg/kg) and xylazine (10 mg/kg) by intraperitoneal injection. Trans-corneal subretinal injections of virus (1 μL) were performed using a 33-gauge blunt needle on a Hamilton syringe. Vector concentrations are detailed within the results section of each experiment. Approximately equal numbers of male and female mice were used. In-life testing and necropsy were performed approximately six-weeks post-injection.

Subretinal injections in cynomolgus monkeys were performed according to previously published methods with minor modification^44^. Two distinct, 75 μL subretinal blebs were created in each eye (one in the superior and one in the inferior retina). Retinotomies were placed extrafoveally with the goal to avoid foveal detachment during surgery. The ability to surgically create multiple, distinct blebs has been previously described^45^. Injected animals received intramuscular methylprednisolone (40 mg) the day prior to surgery and weekly for 4 weeks. Triamcinolone acetonide (8 mg/eye) was administered subconjunctivally to both eyes on the day of surgery.

### In life assessments

Optical coherence tomography (OCT) analysis was performed on anesthetized/dilated mice using the spectral domain OCT system (Bioptigen, Inc., Durham, NC). Post-imaging outer nuclear layer (ONL) measurements were performed with commercial software (InVivoVue, Bioptigen, Inc.) as previously described^46^. Briefly, three regions of the OCT scans were selected for measurement: 3 mm above the center of the optic nerve, the optic nerve meridian, and 3 mm below the center of the optic nerve. A corresponding grid containing eight measurement points, centered on the optic nerve was created using these parameters. At each point, the ONL thickness was measured from the outer plexiform layer to the external limiting membrane using the software’s calipers. OCT data was analyzed by two-way ANOVA with multiple comparisons, using BSS-injected controls as the reference group.

OCT imaging of anesthetized/dilated NHPs was performed using the Heidelberg Spectralis HRA/OCT system. Single, horizontal, high resolution line scans were captured from the optic nerve through the fovea as well as a volume scan that included dosing sites. For the non-GLP study (**Table S3**) imaging was performed at baseline, on Day 1 and at Week 4 and Week 9. Widefield color fundus imaging was performed at the same time points using the Clarity RetCam Shuttle. Ophthalmic examinations were performed by a staff Ophthalmologist via indirect ophthalmoscopy and slit-lamp biomicroscopy at baseline, on Days 2, 7, 14, 28, 56/57, and Week 9.

In the GLP toxicology study (**Table 1**), ophthalmic examinations, intraocular pressure (IOP) measurements, widefield color fundus imaging, OCT imaging, and ERG were performed at pretreatment and at scheduled intervals post-injection through the terminal endpoint (day 183 ± 5 for Groups 1–5; day 92 ± 3 for Group 6). Ophthalmic examinations were performed by a staff veterinary ophthalmologist via indirect ophthalmoscopy and slit-lamp biomicroscopy at pretreatment and on Days 3 (±1), 7 (±1), 14 (±1), 30 (±3), 60 (±3), 90 (±3), 120 (±3), 155 (±3), and prior to terminal necropsy on Day 180 (±3). Anterior and posterior segment findings, including aqueous flare, aqueous cells, vitreous haze, vitreous cells, and retinal perivascular sheathing, were graded using a 0–4 scale per Charles River Laboratories standard operating procedures, modified from the Standardization of Uveitis Nomenclature (SUN) Working Group criteria. Cell color was recorded qualitatively. IOP was measured in both eyes of each animal at the same timepoints as ophthalmic examinations via rebound tonometry. Widefield color fundus imaging was performed at pretreatment and on Day 1 (post-dose) using the Clarity RetCam Shuttle. OCT imaging was performed at pretreatment and on Days 30 (±2), 90 (±3), 120 (±3), 155 (±3), and prior to terminal necropsy using the Heidelberg Spectralis HRA/OCT system; single, horizontal, high-resolution line scans were captured from the optic nerve head through the fovea, as well as a volume scan that included the dosing sites, with images recorded inside and outside the bleb area. For ERG, OCT, and ocular imaging procedures requiring sedation, animals were anesthetized with dexmedetomidine (0.02 mg/kg, IM) and ketamine (12–15 mg/kg, IM). The establishment of the NOAEL was an independent decision made by the study director at the performing CRO.

### Electroretinography

Full-field electroretinography (ERG) was performed on cynomolgus macaques in the GLP toxicology study at pretreatment and on Days 30 (±2), 90 (±3), 120 (±3), 155 (±3), and prior to terminal necropsy. Animals were dark-adapted for a minimum of one hour prior to recording. ERG responses were elicited using a Ganzfeld dome stimulus (Retiport Gamma, Roland Consult) with flash intensities according to ISCEV standard parameters. Scotopic (dark-adapted) responses were recorded followed by photopic (light-adapted) responses following five minutes of light adaptation. Amplitude and latency values were measured from tracings for the a-wave and b-wave. Prior to ERG recording, animals were anesthetized with dexmedetomidine (0.02 mg/kg, IM) and ketamine (12–15 mg/kg, IM) and pupils were dilated with topical proparacaine, phenylephrine, tropicamide, and atropine (administered the day prior). ERG data were analyzed using a repeated measures analysis of variance (RMANOVA) model with PROC GLIMMIX (SAS Institute, Cary, NC). Group, Time, and their interaction were included as fixed factors, with the

Between-Within (BW) degrees of freedom approximation. Data were analyzed assuming a normal distribution of errors with animal within treatment as the R-side random subject repeated across time. The corrected Akaike’s Information Criterion (AICC) was used to select between First-Order Autoregressive and Heterogeneous First-Order Autoregressive covariance structures. If no significant (p > 0.05) Treatment×Time interaction was identified, the Treatment main effect was evaluated. Pairwise comparisons were constructed between vehicle (Group 1) and each treated group (Groups 2, 3, and 4), with results adjusted for multiple comparisons using the method of Edwards and Berry.

### Genome & Transcript Analysis

Eyes from subretinally injected mice were enucleated approximately six-weeks post-injection. Eyes from male and female mice were used at random for genome/transcript analysis. All samples were immediately flash-frozen in RNA*later* (Qiagen, 76104) and stored at ≤-60°C. Both DNA and RNA were extracted from each sample using the Qiagen AllPrep DNA/RNA Mini Kit (Qiagen, 80204) following the manufacturer’s protocol. Following extraction, RNA samples were treated with DNaseI (ThermoFisher, AM2222) to remove residual genomic DNA, and 125 ng of RNA from each sample was reverse transcribed into cDNA (Bio-Rad, iScript cDNA Synthesis Kit, 1708840). An identical aliquot from each sample was not reverse transcribed, to serve as a negative control (-RT). All cDNA and -RT samples were cleaned prior to use to remove remaining buffers and reagents (Zymo, DNA Clean and Concentrator-5, D4004). DNA samples were used directly following extraction.

Samples were analyzed using qPCR or qRT-PCR, respectively using a CFX96 PCR Detection System with C100 Touch Thermal Cycler (BioRad, iQ SYBR Green Supermix, 1708880). AAV-mediated *MYO7A* and endogenous *18S* genome copies/transcript were analyzed using three primer sets: (1) *MYO7A* Front to amplify a 214-bp region on the 5’ end of the front half vector, (2) *MYO7A* Back to amplify a 186-bp regions located at the 3’ end of the back half vector, and (3) *18S* primers that were previously described (**Table S9**)^22^. All primers were confirmed to amplify both vector genomes (DNA) and transcript. Prior to experimental use, all primer sets were validated according to minimum information for publication of quantitative real-time PCR experimental standard^47^. All qPCR and qRT-PCR reactions were performed using the following conditions: (1) initial denaturation at 95°C for 3 minutes, (2) 34 cycles of 95°C denaturation for 30 seconds, 58°C annealing for 30 seconds, 72°C elongation for 1 minute, followed by a plate read step. All samples were run in triplicate and results were determined from the average of the replicate reactions using the ΔΔCq method^48^ with *MYO7A* signal normalized to *18S*. -RT samples were analyzed and verified to be negative for all cDNA samples.

### Western Blot & Jess™ Simple Western

Cells were harvested approximately seventy-two hours post-transduction. Eyes from injected mice were enucleated approximately six weeks post-injection. Retinal punches were collected from injected NHPs in the non-GLP study at approximately nine weeks post-injection. In the GLP toxicology study, retinas from the HA-tagged cohort (Group 6) were collected at approximately three months post-injection (day 92 ± 3) and analyzed by both Jess™ western blot. Four (6 mm in diameter) retinal punches were collected from each primate eye: one punch was taken from the center of each of the two distinct subretinal blebs (bleb 1 and bleb 2), one punch from the macula, and one punch from a distal region of the retina away from the blebs and the macula (superior nasal region). All samples were immediately flash frozen and stored at ≤−60°C until ready for use. All samples were homogenized in RIPA lysis buffer (ThermoFisher, 89901) with added protease inhibitors (ThermoFisher, 78429). Total protein concentration was measured using BCA assays (ThermoFisher, A55864) prior to use.

For traditional western blots, equivalent amounts of total protein were loaded on 8-12% Tris-Acetate gels (Invitrogen, EA0375) and run for approximately 90 minutes at 120V (Invitrogen, A25977). Gels were transferred to a PVDF membrane using a semi-dry system (BioRad, 1704150). The membrane was blocked for 1 hour at room temperature in Licor blocking buffer (927-40000) and incubated at 4°C overnight with primary antibodies (MYO7A: Santa Cruz, sc-74516, 1:500; Vinculin: SantaCruz, sc73614, 1:1000). Secondary antibody (Licor, 926-68072) incubation took place the following day at room temperature for one hour. Membranes were visualized using a Licor Odyssey CLx Imager. Protein quantification was performed by measuring the mean gray value for each sample using the open-source *Fiji* software^49^. Each sample value was normalized to vinculin (VCL) expression.

For capillary electrophoresis analysis, samples were analyzed using Jess™ system (ProteinSimple, Bio-Techne) as previously described^50^. For all studies, samples were analyzed using the standard protocol for the 66-440 kDa separation module (Bio-Techne, SM-W008). For cell lysate and mouse eye samples, lysates were diluted to 2.0 µg/uL in 0.1X Sample Buffer followed by an incubation at 100°C for 10 minutes. The primary antibodies, anti-MYO7A (SantaCruz, sc-74516, 1:5) or anti-VCL (Cell Signaling Technologies, 13901, 1:50), were diluted in Antibody Diluent 2. MYO7A was normalized to VCL using the Replex module (Bio-Techne, RP-001) as detailed in (Calabro et al., 2023). For NHP samples, lysates were diluted to 0.5 µg/uL in 0.1X Sample Buffer followed by an incubation at 100°C for 10 minutes. The primary antibodies, anti-MYO7A (SantaCruz, sc-74516, 1:50) or anti-HA (Cell Signaling Technologies, 3724, 1:200), were diluted in Antibody Diluent 2. Samples were normalized using the Total Protein Normalization module (Bio-Techne, DM-PN02).

Of note, apparent molecular weights may differ slightly between Jess capillary electrophoresis and traditional SDS-PAGE western blots due to differences in separation matrix. An example of this phenomenon can be seen when comparing the size of full length MYO7A in the traditional western blot in panel D of Figure 1 (∼250 kDa) vs. all subsequent figures (∼225 kDa).

### Immunohistochemistry

Mice were sacrificed six weeks post-injection. Eyes were enucleated and fixed in 4% paraformaldehyde (PFA) overnight, and eyecups were dissected and prepared for sectioning according to established methods^51^. Eyes were oriented vertically in optimal cutting temperature medium (Sakura, 4583) and flash frozen. Cryosections (10 µm) were cut and fixed to microscope slides using a Leica CM3050 S cryostat. Sections were permeabilized using 5% Triton X-100 detergent in PBS and blocked using 1% bovine serum albumin (BSA) in PBS. Cryosections were incubated overnight at 4°C with anti-hemagglutinin (HA) rabbit monoclonal antibody (Cell Signaling Technology, 3724) diluted 1:800 in PBS containing 0.3% Triton X-100 and 1% BSA. Immunoglobulin G (IgG) secondary antibody Alexa Fluor 488 (Thermo Fisher, Z25302) diluted 1:500 in PBS was added for one hour at room temperature. Sections were counterstained with DAPI diluted 1:10,000 for five minutes at room temperature. Images were taken using a Leica TCS SP8 confocal microscope. Gain and exposure settings remained constant for all images.

Whole macaque eyes were enucleated and perfused with approximately 100 µL of 4% PFA. A 6 mm hole was created in the cornea to facilitate the fixation process. The whole eye was then placed in 4% PFA overnight. The next day, eyes were transferred and stored in 0.1M PBS/0.03% sodium azide solution until ready for further processing. Eyes were dissected, and the eye cup was successively incubated in 10% sucrose for 30 minutes, 20% sucrose for 30 minutes, and 30% sucrose overnight at 4°C. The following day, eyes were placed in a 3:2 mixture of 30% sucrose/embedding media (Sakura, 4583). Then, eyes were embedded, flash frozen, and stored at ≤-60°C. Sections were cut (12µm) and transferred to glass slides. Cryosections were incubated with 0.5% Triton X-100 for 1 hour and then blocked for 1 hour with 10% normal goat serum (ThermoFisher, 50062Z). Sections were incubated with anti-HA antibody at 1:800 overnight at 4°C. The next day, anti-rabbit Alexa Fluor 555 (dilution 1:500, ThermoFisher, A32732) in 10% normal goat serum was added to the slides for 90 minutes at room temperature, followed by counterstaining with DAPI. Slides were then incubated for 5 minutes with Autofluorescence Eliminator (Millipore Sigma, 2160). Images were acquired using a Leica TCS SP8 confocal microscope or Keyence fluorescent microscope.

## DATA AVAILABILITY STATEMENT

The authors confirm that the data supporting the findings of this study are available within the article and/or its supplemental information.

## ACKNOWLEDGEMENTS

This work was funded by sponsored research agreements from Atsena Therapeutics (to SEB). We would like to thank Lauren Knupp, McKenzie Skiendziel and Baylee Tucker at Charles River Laboratories for their contributions to our non-GLP NHP study and Mike Kelly for oversight of the manufacturing of the ATSN-301 tox lot vector.

## AUTHOR CONTRIBUTIONS

Conceptualization (KRC, SLB, EAM, SEB); Funding Acquisition (SLB, SEB); Writing original draft (KRC, SEB); Writing-review and editing (CK, PSN, SLB, SEB); Data Curation (KRC, SLB, CK, EMX, SMC, DF, HLF, JJP, WL, HZ, VS, HR, ML, RFB); Formal Analysis (KRC, SLB, CK, HR, RFB, EAM, SEB).

---

**Figure S1.**
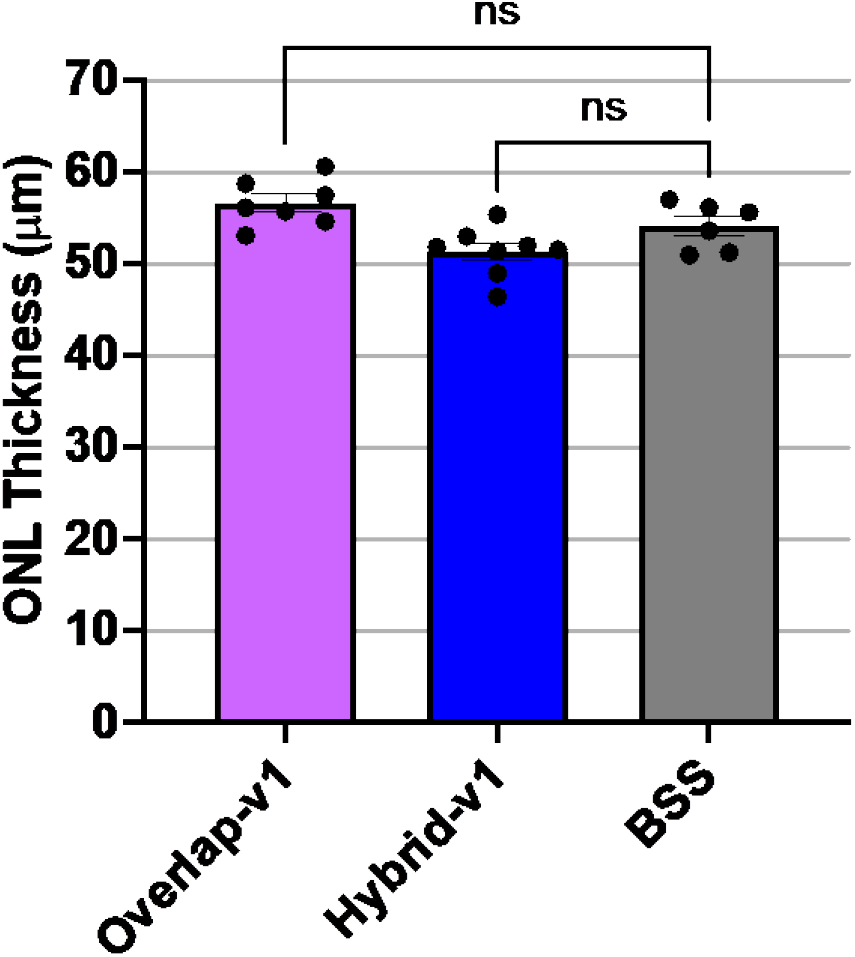
Analysis of retinal structure in *Myo7a* KO mice subretinally injected with first-generation dual AAV-MYO7A vectors. Average outer nuclear layer (ONL) thickness measured by optical coherence tomography (OCT) in *Myo7a* KO mice subretinally injected with overlap-v1 or hybrid-v1 dual AAV8(Y733F)-MYO7A vectors or balanced salt solution (BSS) vehicle (n = 6–8 eyes per treatment) at six weeks post-injection. No statistically significant differences in ONL thickness were observed between vector-injected and BSS-injected groups. ONL = outer nuclear layer; BSS = balanced salt solution/vehicle injection; ns = not statistically significant. Statistics performed by two-way ANOVA with multiple comparisons using BSS as the control group.

**Figure S2.**
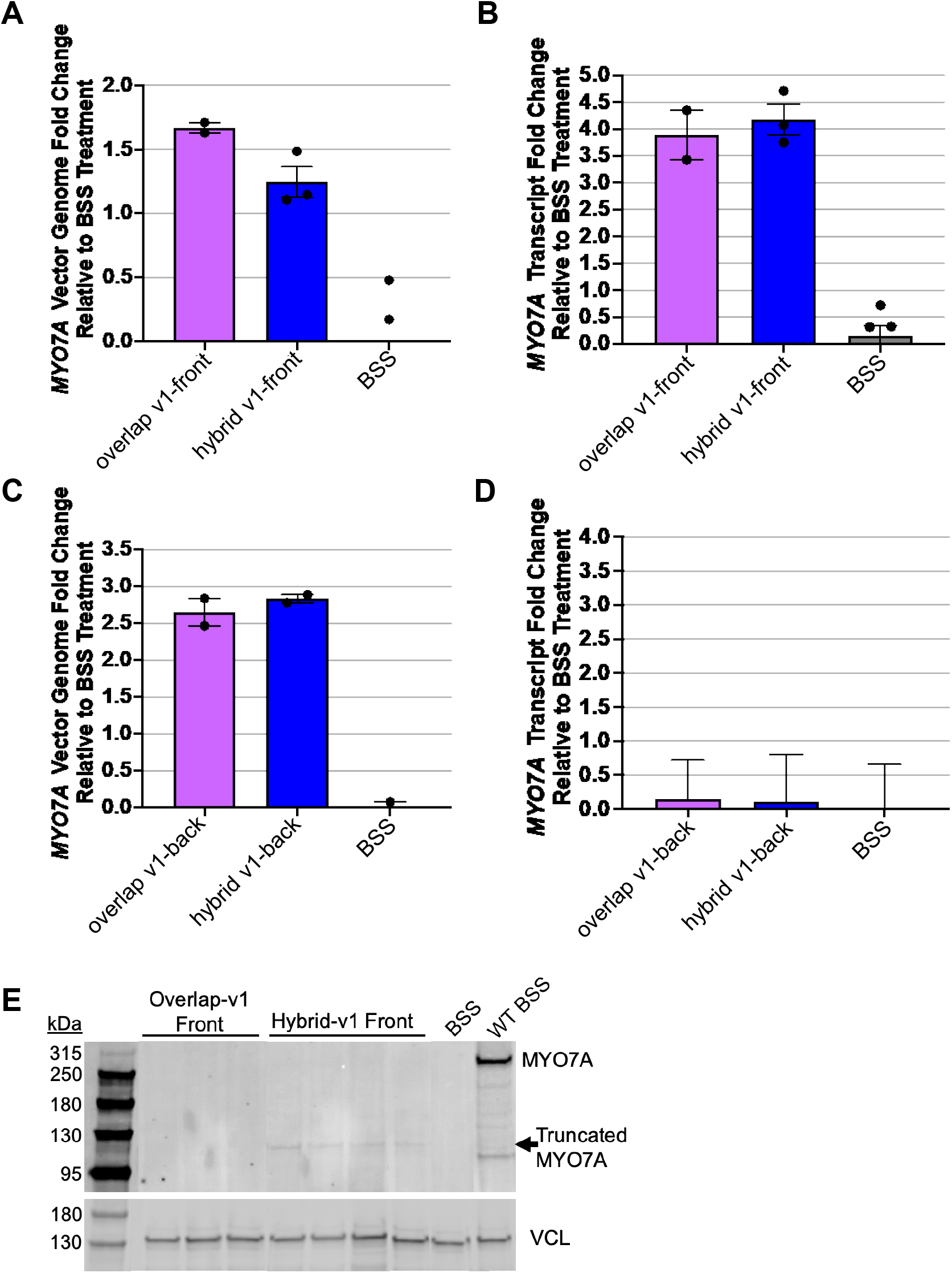
Analysis of vector genomes, transcripts, and protein in *Myo7a* KO mouse eyes injected with front or back half vector alone. (A) *MYO7A* vector genome copies detected by qPCR in *Myo7a* KO mouse eyes injected with front half overlap-v1 or hybrid-v1 vector alone (n = 2–3 eyes per treatment), normalized to *18S* genomic DNA and expressed as fold change relative to BSS-injected controls. Front half vector genomes were detectable in all samples. (B) *MYO7A* transcript levels detected by qRT-PCR in *Myo7a* KO mouse eyes injected with front half overlap-v1 or hybrid-v1 vector alone, normalized to *18S* transcript and expressed as fold change relative to BSS-injected controls. Transcript was detected in all samples, with comparable levels between platforms. (C) *MYO7A* vector genome copies detected by qPCR in *Myo7a* KO mouse eyes injected with back half overlap-v1 or hybrid-v1 vector alone (n = 2–3 eyes per treatment), normalized to *18S* genomic DNA and expressed as fold change relative to BSS-injected controls. Back half vector genomes were detectable in all samples. (D) *MYO7A* transcript levels detected by qRT-PCR in *Myo7a* KO mouse eyes injected with back half overlap-v1 or hybrid-v1 vector alone, normalized to *18S* transcript and expressed as fold change relative to BSS-injected controls. Transcript levels were negligible regardless of platform. (E) Western blot of retinal lysates from *Myo7a* KO eyes injected with front half overlap-v1 or hybrid-v1 vector alone (n = 3–4 eyes per treatment). Truncated MYO7A (∼116 kDa) was detectable only in eyes injected with hybrid-v1 front half vector, corresponding to the protein expected to be translated from the promoter-containing front half vector. No truncated protein was detected in eyes injected with overlap-v1 front half vector. BSS = balanced salt solution/vehicle-injected Myo7a KO eyes; WT BSS = balanced salt solution/vehicle-injected C57BL/6J eyes; kDa = kilodaltons; VCL = Vinculin.

**Table S1:**
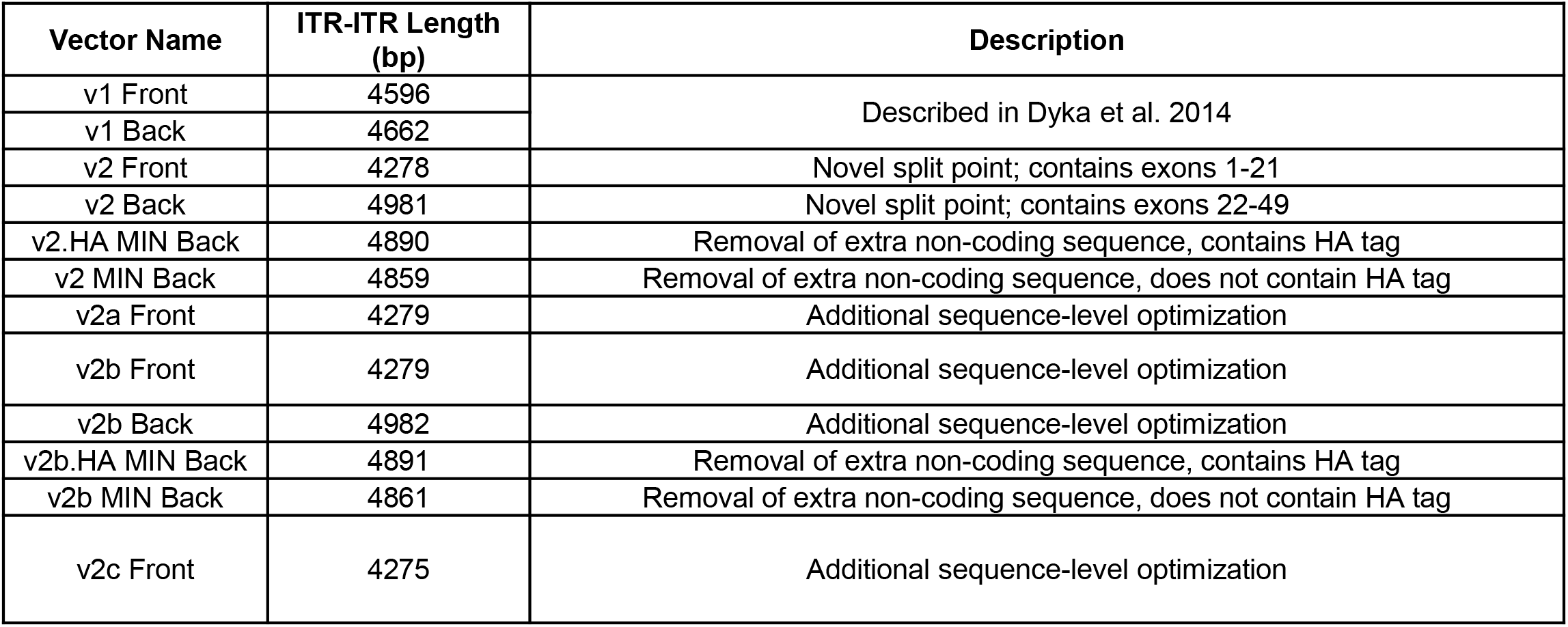
Summary of Hybrid Vector/Modifications.

**Table S2.**
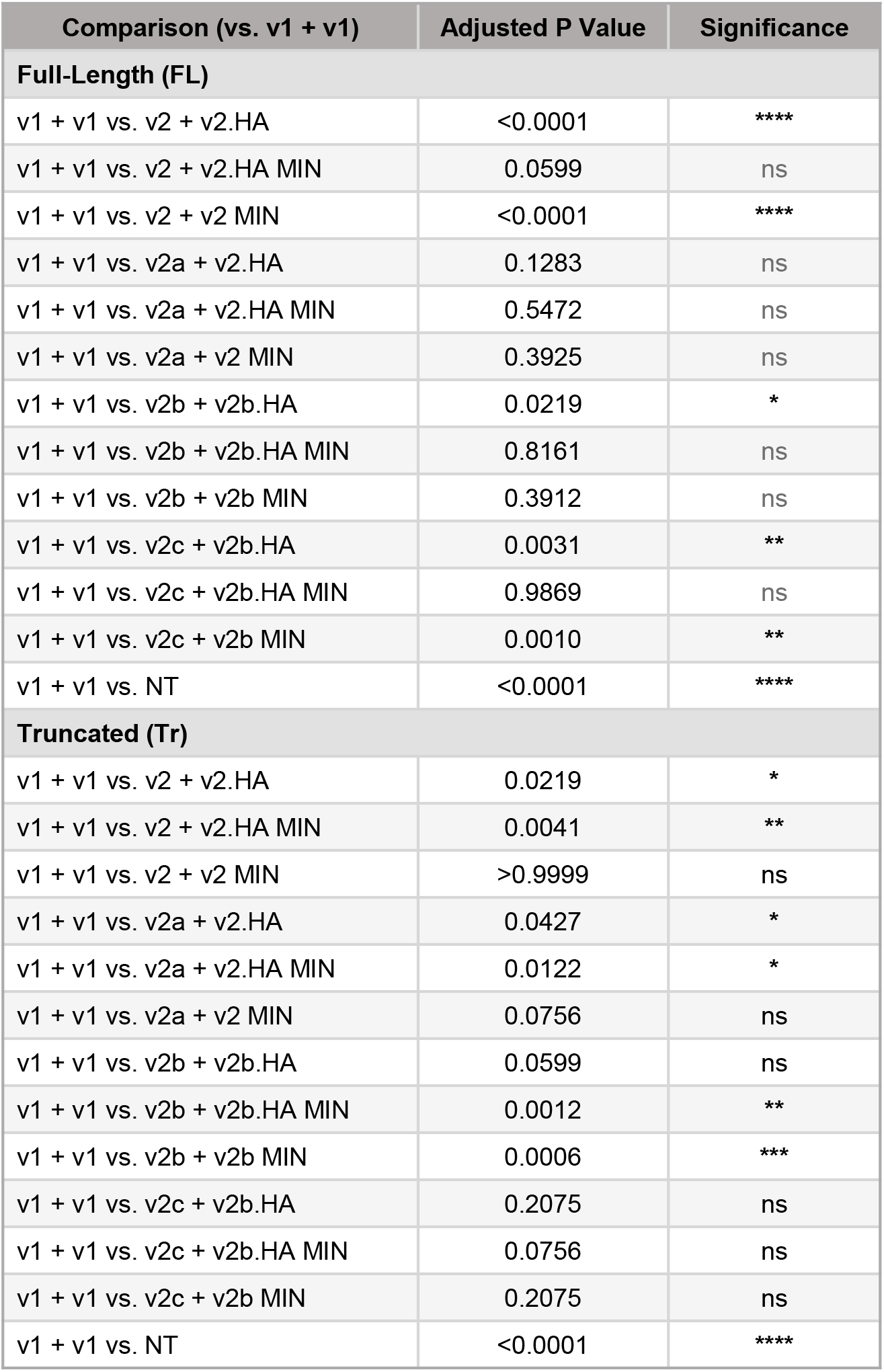
Summary of Overlap Vectors/Modifications. Statistical comparisons for Full Length (FL) and Truncated (Tr) expression. Two-way ANOVA with Dunnett’s multiple comparisons test. Each construct combination was compared to the v1 + v1 reference within its own FL or Tr group (13 comparisons per family; family-wise α = 0.05). Data correspond to the bar graph in Figure 2. ns = not significant. *P < 0.05; **P < 0.01; ***P < 0.001; ****P < 0.0001; ns, not significant (P ≥ 0.05).

**Table S3.**
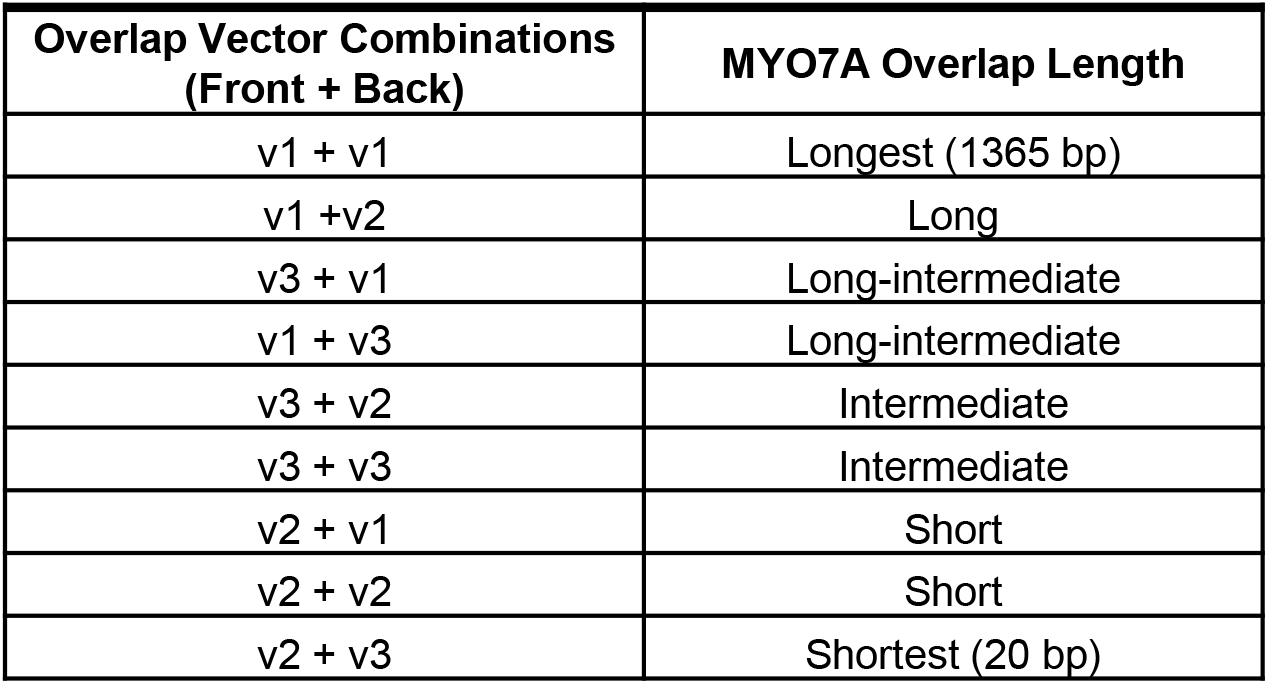
Summary of Overlap Vectors/Modifications.

**Figure S3.**
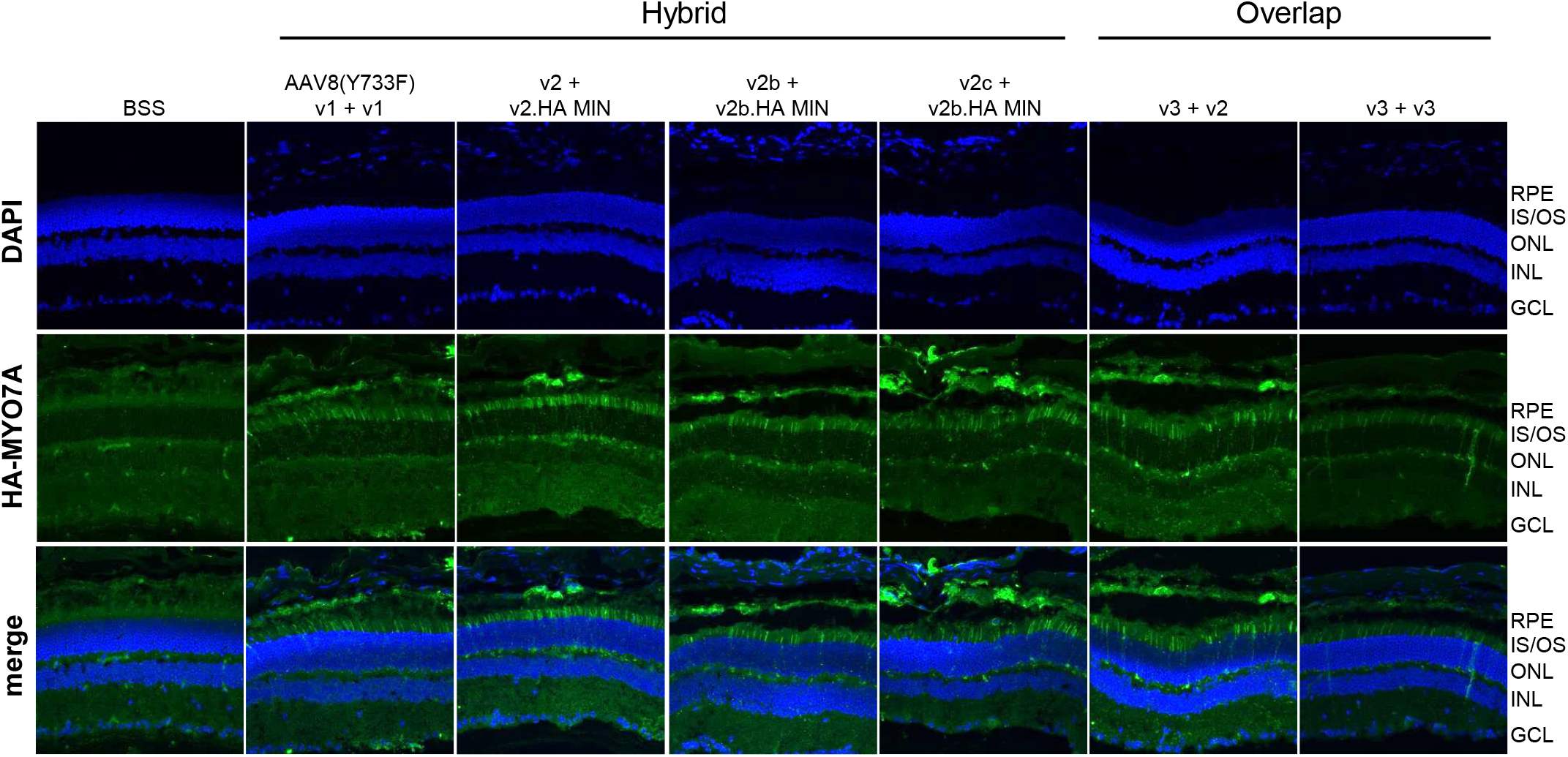
Immunohistochemical analysis of retinal sections from *Myo7a* KO mice subretinally injected with optimized hybrid and overlap dual AAV.MYO7A vector pairs. Cryosections from *Myo7a* KO mouse eyes subretinally injected with BSS vehicle, AAV8(Y733F) hybrid v1+v1, AAV.SPR hybrid v2+v2.HA MIN, AAV.SPR hybrid v2b+v2b.HA MIN, AAV.SPR hybrid v2c+v2b.HA MIN, AAV.SPR overlap v3+v2, or AAV.SPR overlap v3+v3 were stained with an anti-HA antibody (green) to detect vector-mediated MYO7A-HA expression and counterstained with DAPI (blue) to label nuclei. Representative images are shown for DAPI (top row), HA-MYO7A (middle row), and merged channels (bottom row). HA-MYO7A signal is localized to the photoreceptor inner and outer segment (IS/OS) layer and retinal pigment epithelium (RPE) across all vector-treated groups. Animals were evaluated at six weeks post-injection. RPE = retinal pigment epithelium; IS/OS = inner segment/outer segment; ONL = outer nuclear layer; INL = inner nuclear layer; BSS = balanced salt solution/vehicle injection; HA = hemagglutinin.

**Figure S4.**
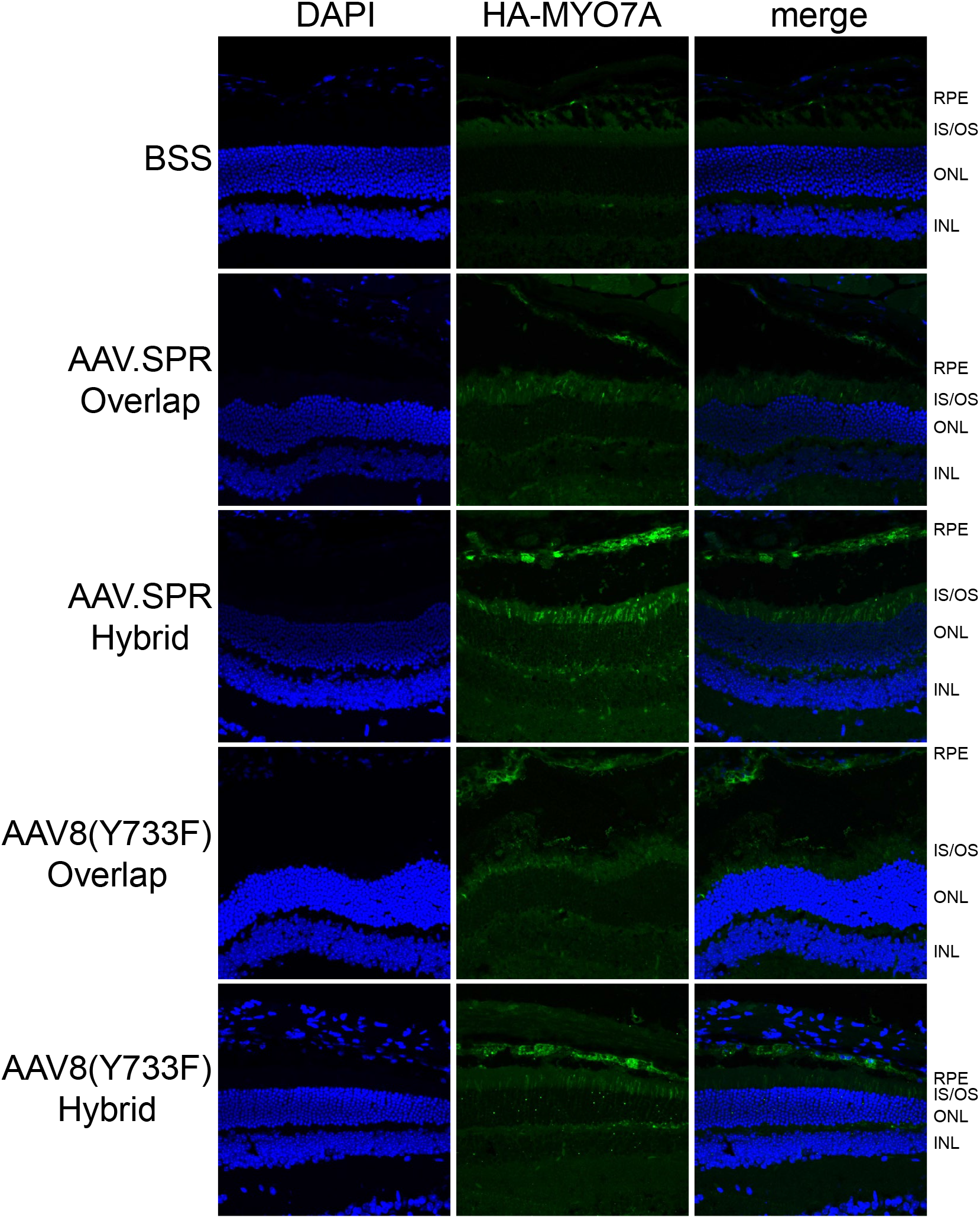
Immunohistochemical analysis of retinal sections from *Myo7a* KO mice subretinally injected with top-performing hybrid and overlap dual AAV.*MYO7A* vector pairs packaged in AAV.SPR or AAV8(Y733F). Cryosections from *Myo7a* KO mouse eyes subretinally injected with BSS vehicle, AAV.SPR overlap (v3+v3), AAV.SPR hybrid (v2c+v2b.HA MIN), AAV8(Y733F) overlap (v3+v3), or AAV8(Y733F) hybrid (v2c+v2b.HA MIN) were stained with an anti-HA antibody (green) to detect vector-mediated MYO7A-HA expression and counterstained with DAPI (blue) to label nuclei. Representative images are shown for DAPI (left column), HA-MYO7A (middle column), and merged channels (right column) for each treatment group. HA-MYO7A signal is localized to the photoreceptor inner and outer segment (IS/OS) layer and retinal pigment epithelium (RPE) in all vector-treated groups, with visibly greater signal intensity observed in AAV.SPR hybrid-injected eyes relative to all other groups. Animals were evaluated at six weeks post-injection. RPE = retinal pigment epithelium; IS/OS = inner segment/outer segment; ONL = outer nuclear layer; INL = inner nuclear layer; BSS = balanced salt solution/vehicle injection; HA = hemagglutinin.

**Table S4.**
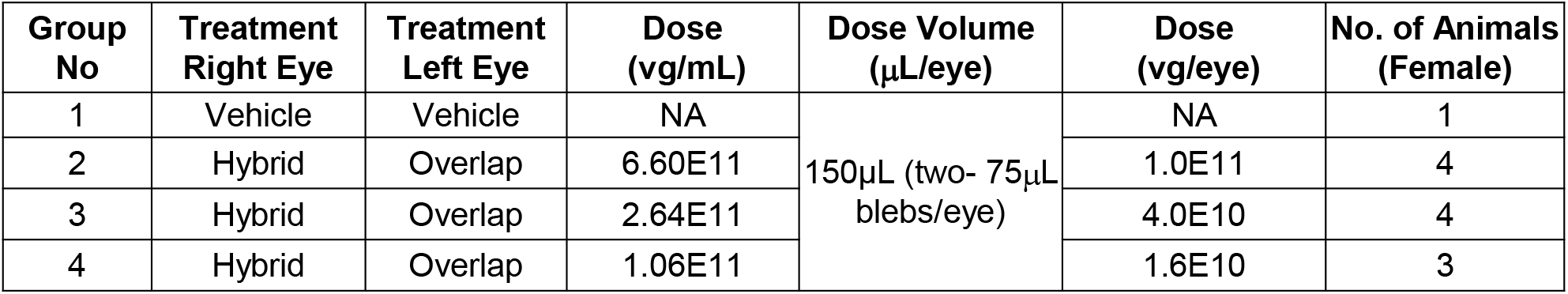
Experimental Design for Testing Lead Hybrid/Overlap sets in NHP.

**Table S5.**
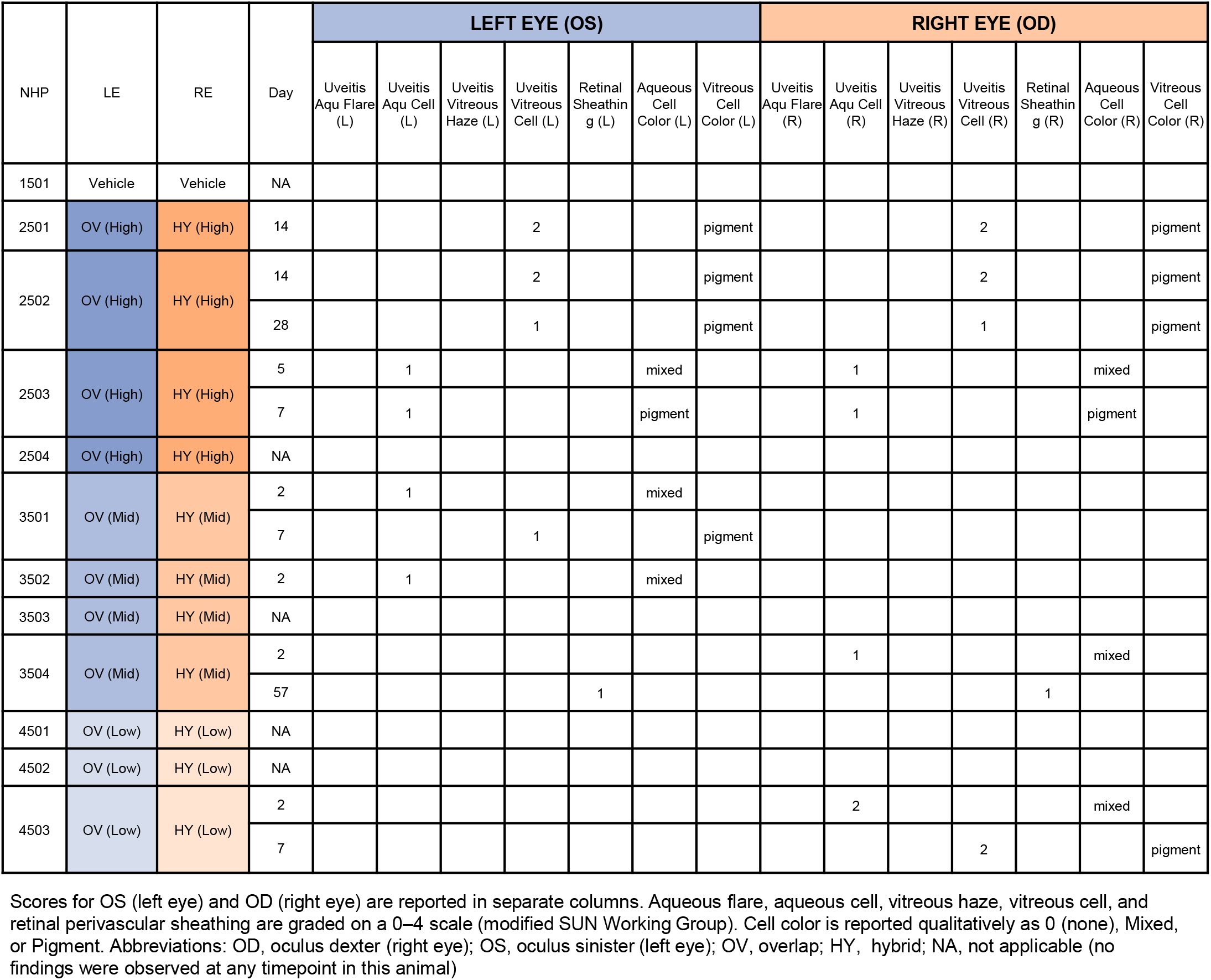
Summary of ophthalmic exams from non-GLP study in NHP.

**Figure S5.**
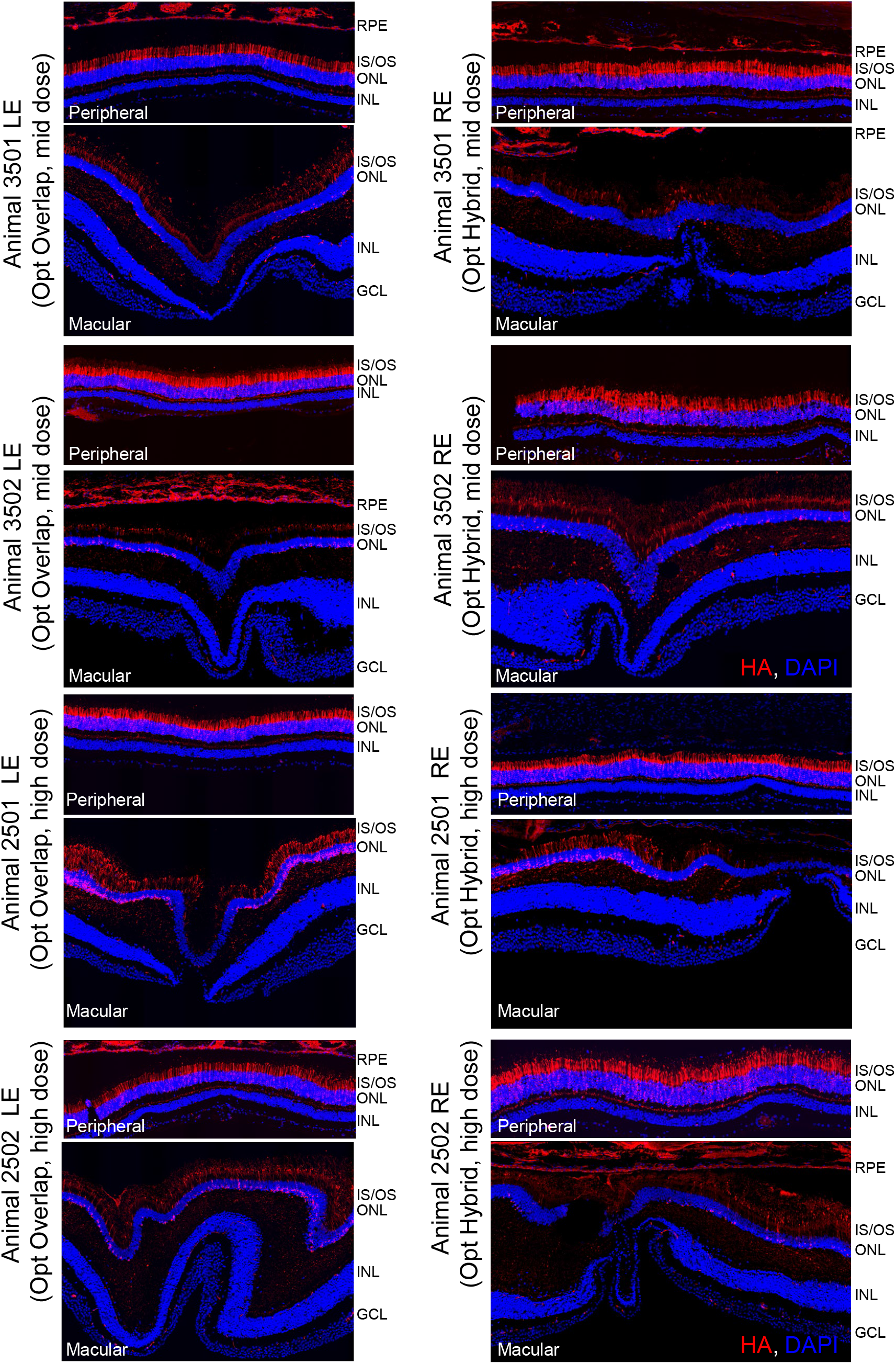
Immunohistochemical analysis of retinal sections from cynomolgus monkeys subretinally injected with AAV.SPR Opt Hybrid or AAV.SPR Opt Overlap dual AAV·MYO7A-HA vectors at mid and high doses. Retinal cryosections from cynomolgus monkeys subretinally injected with mid dose (3.9E10 vg/eye; Animals 3501 and 3502) or high dose (9.9E10 vg/eye; Animals 2501 and 2502) of AAV.SPR Opt Hybrid (right eye) or AAV.SPR Opt Overlap (left eye) dual vectors were co-stained with anti-HA (red) to detect vector-mediated HA-tagged MYO7A expression and counterstained with DAPI (blue) to label nuclei. Representative images are shown for peripheral retina (within the subretinal injection bleb) and macula for each animal and treatment group. HA-tagged MYO7A signal is localized to the photoreceptor inner and outer segments (IS/OS) and retinal pigment epithelium (RPE) in peripheral and macular retinal sections from animals receiving both platforms at both doses, with expression in the macula present at lower levels relative to the bleb region. Animals were evaluated at approximately nine weeks post-injection. RE = right eye; LE = left eye; ONL = outer nuclear layer; INL = inner nuclear layer; IS/OS = inner segment/outer segment; RPE = retinal pigment epithelium; GCL = ganglion cell layer; HA = hemagglutinin; DAPI = 4′,6-diamidino-2-phenylindole

**Table S6.**
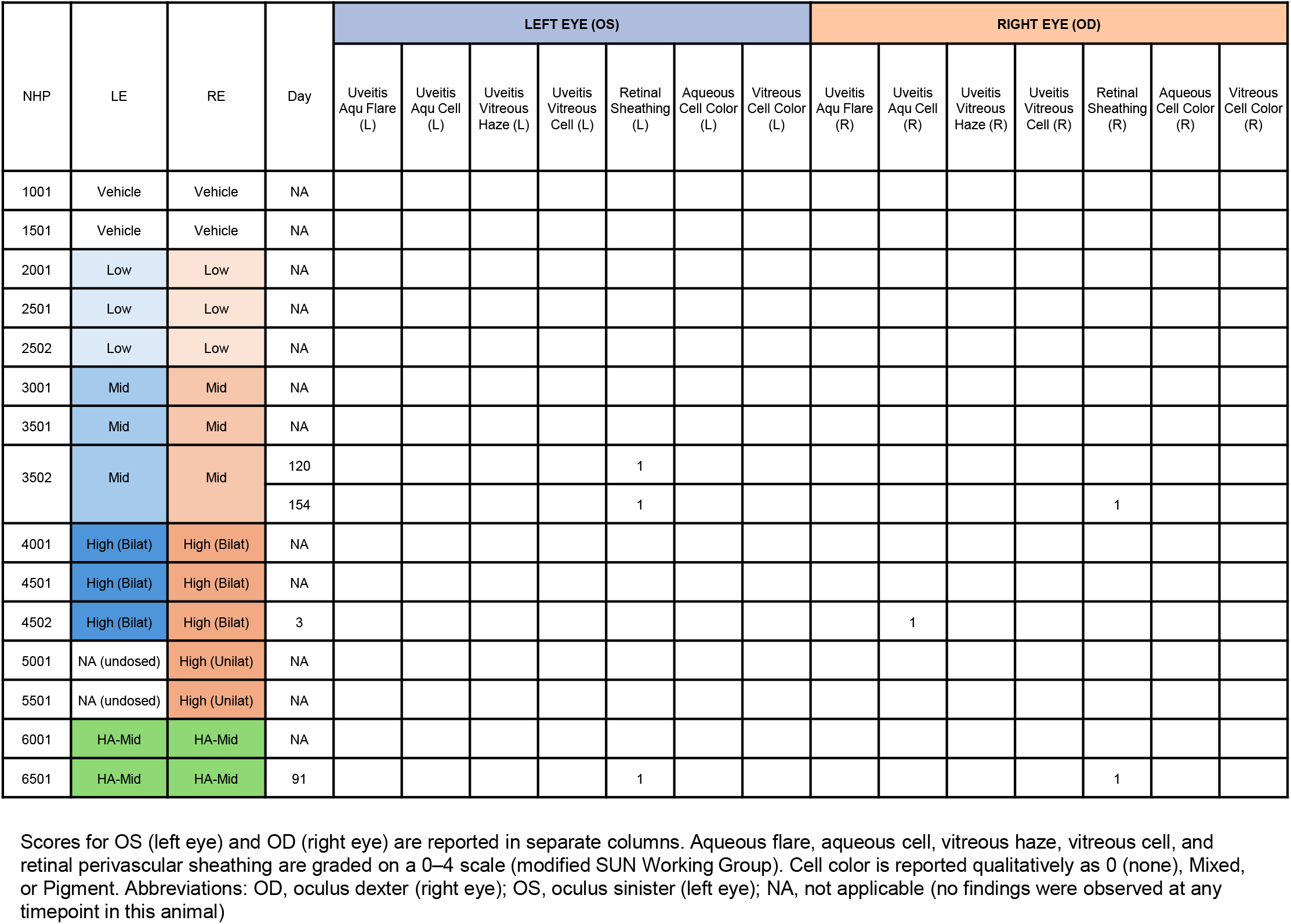
Summary of ophthalmic examination findings from the 6-month GLP toxicology study in cynomolgus macaques administered ATSN-301 by subretinal injection.

**Table S7.**
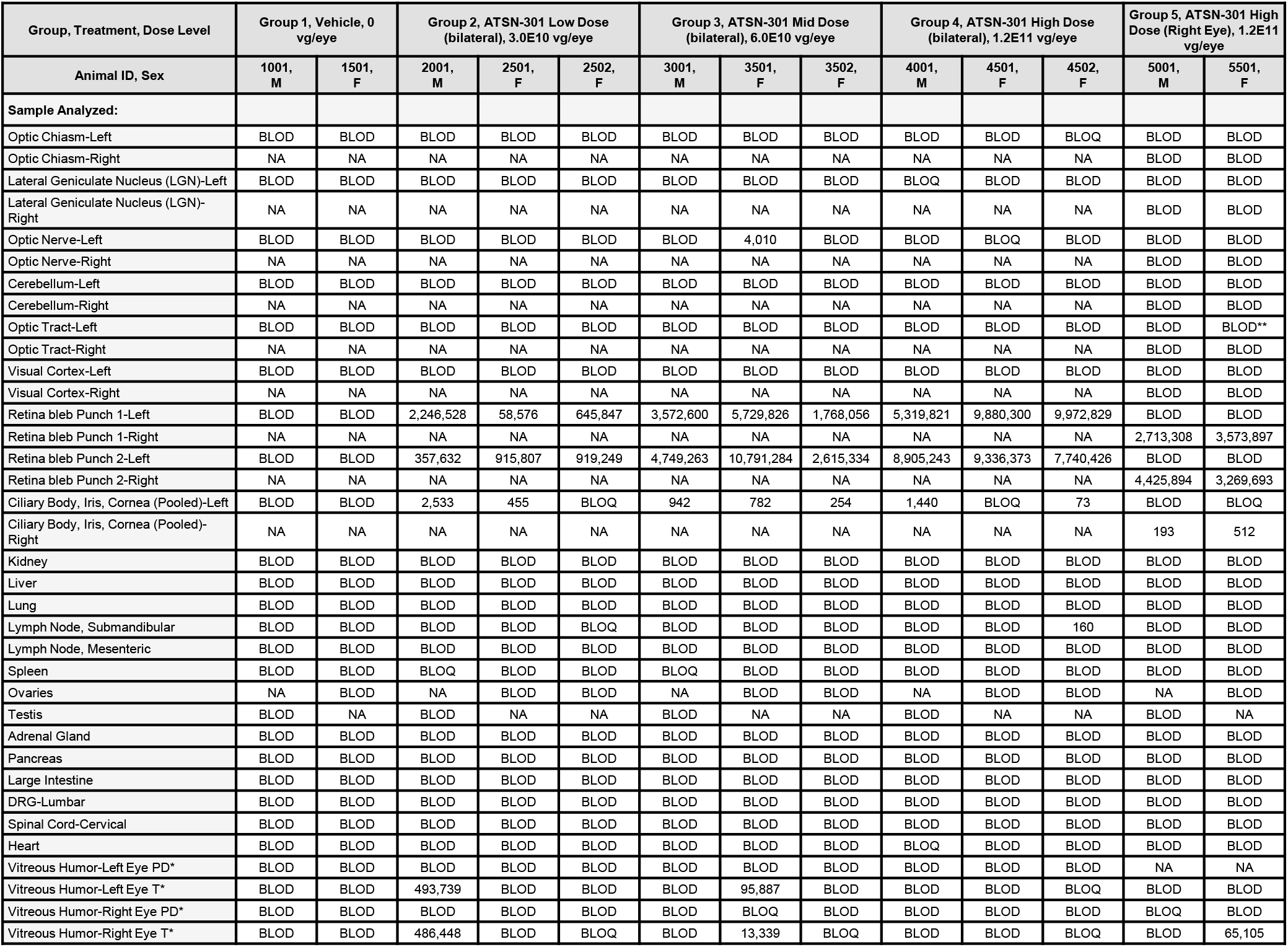
Individual Tissue Biodistribution Data (ds vg per µg of DNA) – ATSN-301-F. A: Qty %CV > 25%, B: Vehicle Control > BLOQ. *Reported as ss copies per mL of humor. **Reported as ds vg per 10 µL of DNA. NA, Not Applicable. BLOD, below limit of detection. BLOQ, below limit of quantification.

**Table S8.**
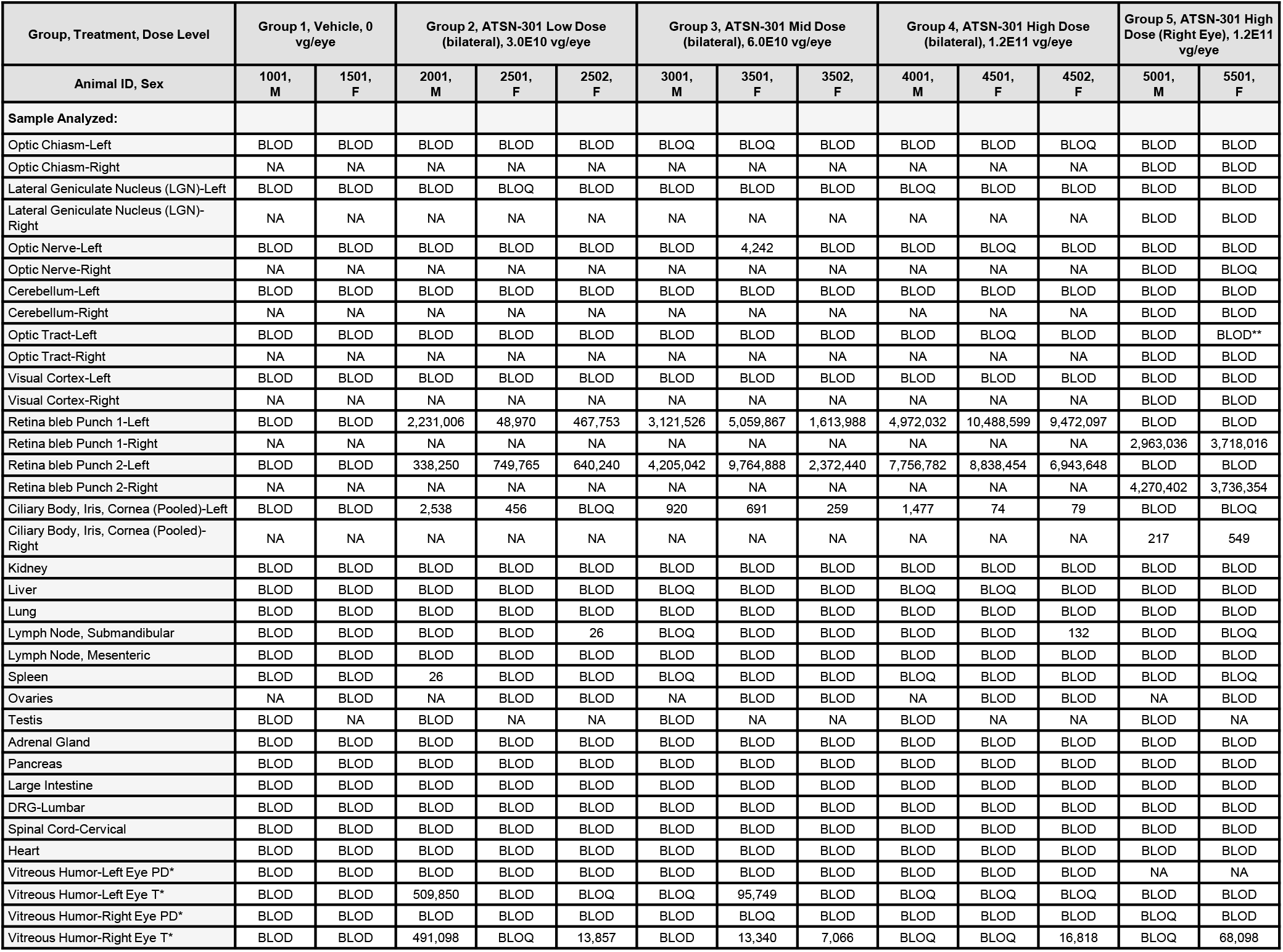
Individual Tissue Biodistribution Data (ds vg per µg of DNA) – ATSN-301-B. *Reported as ss copies per mL of humor. **Reported as ds vg per 10 µL of DNA. NA, Not Applicable. BLOD, below limit of detection. BLOQ, below limit of quantification.

**Table S9.**
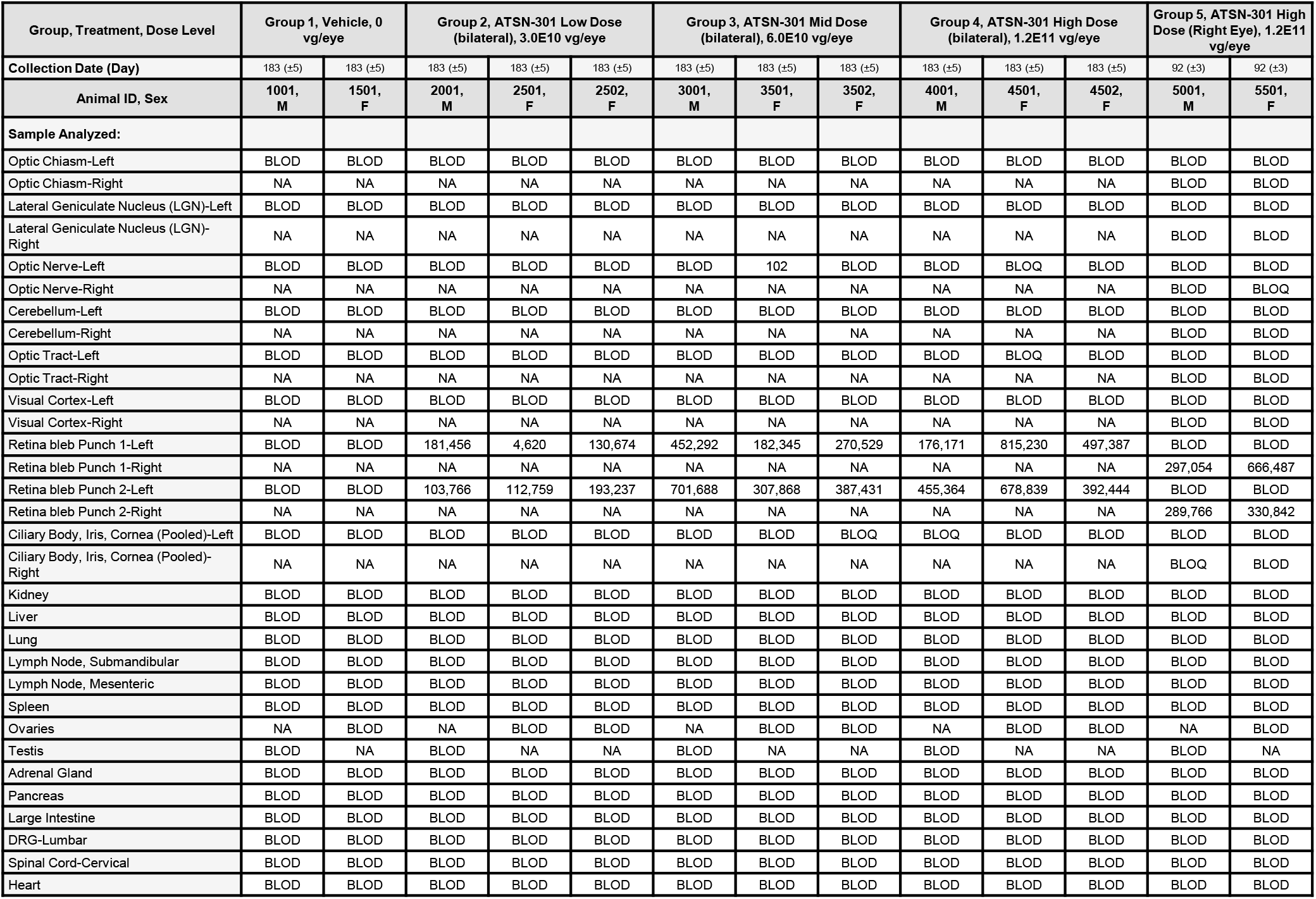
Individual Tissue ATSN-301 Vector-derived mRNA Data (ss copies per 100 ng of total RNA) NA, Not Applicable, Not Analyzed. BLOD, below limit of detection. BLOQ, below limit of quantification.

**Table S10.**
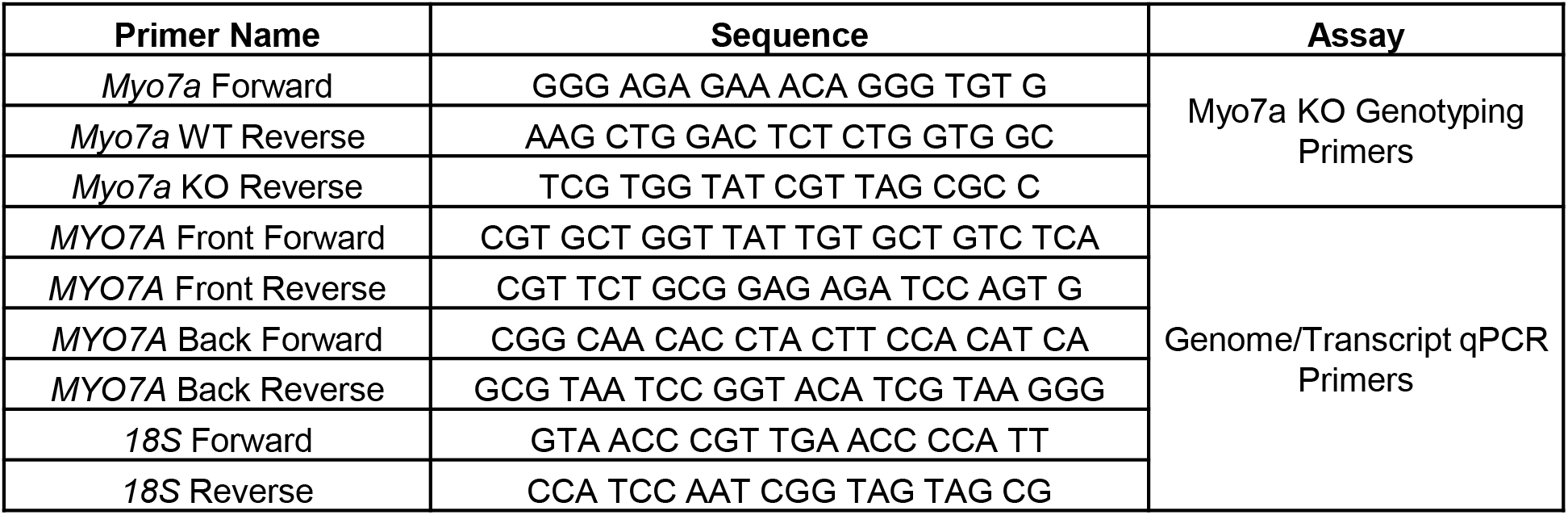
List of primers used to amplify murine *Myo7a* or human *MYO7A* and 18S internal control.

